# Prediction and principle discovery of drug combination based on multimodal friendship features

**DOI:** 10.1101/2024.08.28.610203

**Authors:** He-Gang Chen, Xionghui Zhou

## Abstract

Combination therapy, which can improve therapeutic efficacy and reduce side effects, plays an important role in the treatment of multiple complex diseases. Yet, the design principles of molecular combinations remain unclear. In addition, the huge search space of candidate drug combinations and the numerous heterogeneous data has brought us a big challenge. Here, we proposed a Friendship based Method (FSM), which integrates diverse drug-to-drug information to predict drug combinations for specific diseases. By quantifying the friendship-based relationship between drugs, we found that there is a moderate similarity between the drugs of effective drug combinations in a high-dimensional, heterogeneous feature space. Following this discovery, FSM applied a two-step strategy to predict clinically efficacious drug combinations for specific diseases. First, our method employs the friendship features to evaluate whether each drug is combinable. Then, the synergistic potential of combinable drugs was further evaluated. FSM was validated on two types of disease. The results show that FSM achieves substantial performance improvement over other state-of-the-art methods and tends to have low toxicity. These results indicate that our model could potentially offer a generic, powerful strategy to identify efficacious combination therapies in the vast search space.

## 1 Introduction

Complex diseases such as cancer and cardiovascular diseases are often caused by variations in multiple signaling pathways or molecular sub-networks, limiting the effectiveness of monotherapy [1]. Combination therapy, with its potential for increased efficacy and reduced individual dosage-related adverse effects, has emerged as a promising treatment strategy [2,3]. Traditionally, the discovery of drug combinations relied on extensive biological experiments and screening [4–6]. However, the vast number of possible combinations among the thousands of FDA-approved drugs and hundreds of potential diseases hinders the identification of effective combinations [7]. Furthermore, the underlying mechanisms of drug combinations remain unclear [8]. Therefore, there is an urgent need for innovative methods to predict effective drug combinations for diseases and to gain insights into the behavior of compound combinations in biological systems, ultimately advancing the development of multicomponent therapies.

Computational approaches hold great promise in streamlining the identification of effective drug combinations [9]. Several successful models have been developed, utilizing post-treatment gene expression changes and drug target information to predict such combinations. For instance, Sun et al. [8] integrated targeting network features and transcriptomic profiles to establish a Ranking-system of Anti-Cancer Synergy. The DrugComboRanker platform was created to prioritize combinations that target alternative and complementary disease signaling modules [10]. Cheng et al. [11] recently demonstrated that statistically significant drug combination efficacy is observed mainly when drug pairs target closely related disease modules and exhibit a complementary relationship. Furthermore, Huang et al. [12] used clinical drug side effect data to predict potential new drug pairs under the assumption that co-prescribed drugs tend to avoid serious adverse reactions. These methods indeed outperform random guessing in prediction performance. However, they primarily rely on a single omics data type, potentially missing crucial information on complex biological mechanisms. Additionally, single-data-type approaches are more susceptible to data-specific noise and may have limited utility and accuracy [13,14].

Data integration approaches have shown promise in enhancing analytical power [13]. With the increasing accessibility of diverse data types [15], novel strategies have emerged for the integration of multiple data types. For instance, Zhao et al. [16] condensed n-dimensional drug pair vectors into one-dimensional similarity values and combined these values into a new k-dimensional feature vector, where k represents the number of data types. Shi et al. [17] developed a two-layer classifier to integrate different drug data types and mitigate the high-dimensional feature vectors generated by feature cascading. Li et al. [18] integrated similarity-based multi-feature drug data using neighbor recommender methods and ensemble learning algorithms to detect unknown drug combinations. Liang et al. [19] applied a node embedding approach to multilayer networks and used it for predicting drug combinations with multilayer graph data. However, these approaches have followed limitations: (1) Some models integrate a few data types; (2) Most methods overlook drug relationships; (3) The current methods lack adequate pre-screening capabilities in the vast search space; (4) Our exploration of combination therapy is still in its early stages, but most methods assume unknown drug pairs are negative samples.

To address these limitations, we introduce a novel feature called “Friendship” in this manuscript. Friendship not only integrates diverse information from heterogeneous data but also enhances drug characterization by incorporating similarity information between each drug and all other drugs. With this feature, we quantify drug-drug relationships and identify effective drug combination patterns for specific diseases within high-dimensional, heterogeneous feature spaces. Building upon this discovery, we present FSM, a new framework for identifying clinically effective drug combinations tailored to specific diseases. To eliminate bias arising from labeling unknown samples as negative, we employ one-class support vector machines (OCSVM) as the classifier in FSM. Comprehensive testing demonstrates that FSM significantly outperforms other state-of-the-art methods. Moreover, results reveal that drug combinations predicted by FSM significantly enrich effective combinations and tend to have low toxicity. These findings affirm FSM as a valuable tool for predicting drug combinations in a vast combinatorial space and shedding new light on combination therapy mechanisms.

## 2 Materials and Methods

### 2.1 Datasets

In this study, we gathered data related to drug targets [11], adverse effects [20], chemical structures [21], pathways (http://www.gsea-msigdb.org/gsea/index.jsp), and clinical information [21] for comprehensive drug characterization. We utilized downloaded data on gold standard drug combinations [11] and adverse drug-drug interactions [21] to assess our model’s performance. For additional information about these datasets, please refer to Supplementary Note 1.

### 2.2 Similarity scores calculation

Chemical Structures similarity: We using a chemical molecular information processing package (Rdkit) implemented in Python to compute MACCS fingerprints of each drug and used the Tanimoto coefficient to calculate the structural similarity between two drugs. If the MACCS fingerprints of two drugs are A and B respectively, the Tanimoto coefficient of the drug-drug pair is defined as:

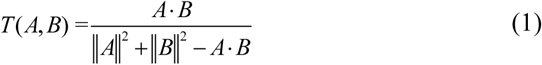

Adverse effects/Target similarity: A Tanimoto correlation was used to measure the degree of similarity for the profiles of two drugs.

Clinical similarity: The drug clinical similarity (*S_atc_*) of drugs A and B is defined via the ATC codes as below:

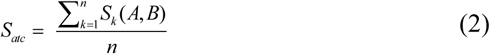

where n represents the level of ATC codes (n = 5 here). A score *S_k_* (*A*, *B*) is used to define the kth level clinical similarity between drugs A and B (notable, drugs can have multiple ATC codes):

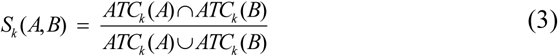

where ATC_k_ represents all ATC codes of a drug at the kth level.

Pathway similarity: The pathway features of drugs are calculated by the drug target and the enrichment degree of pathways. We used a permutation test (Supplementary Note 2) to calculate the enrichment of drug targets and each pathway. If drugs were enriched, the score is 1 (*P*-value ≤ 0.01). Otherwise, it is 0. Finally, we got the 186-dimensional pathway feature for each drug and then used the Tanimoto coefficient to measure the similarity of pathways between different drugs.

### 2.3 Friendship feature construction

The Friendship feature of a drug consists of the interaction between the drug and all the other drugs. Given a set of *n* drugs, denoted as *D = {d_1_, d_2,_…, dn} ⊂R^m^,* The Friendship feature of di is defined as below.

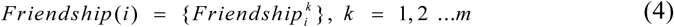

Where m represents the number of data types. The *Friendship^k^* represents the Friend features at the k_th_ level, which was defined as follow:

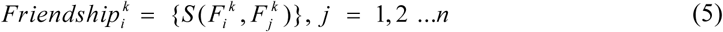

Where n represents the drug number and *S* (*F ^k^*, *F ^k^*) indicates that the similarity between the drug feature vectors at the kth level. Let *Friendship*(*i*, *j*) is the friendship feature of combinatorial drug *d_i_* and *d_j_*. Considering the symmetry of the combinatorial drug (*Friendship(i, j) = Friendship (j, i)*), we defined *Friendship*(*i*, *j*) as follows:

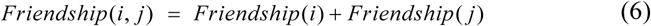

### 2.4 The threshold for Combinatoriality of drugs

Youden’s index γ [22] has been used to evaluate tests diagnostic [23]. It’s derived from sensitivity and specificity and denotes a linear correspondence balanced accuracy.

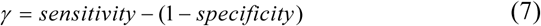

In our work, γ is used as a measure to identify the optimal threshold for drug combinatoriality. In the training data set, we used 100 times two-cross validation to calculate the best cut-off. In each round of sampling, we cross-validated all monotherapies in the training set and found the threshold corresponding to the maximum value of γ. Finally, we took the average value of 200 thresholds as the final threshold in the model, which evaluates whether each drug is combinatorial.

### 2.5 Positive drug combinations selection

Through 2-fold cross-validation, 5% drug combination with the highest score in the test set is selected as the candidate drug pairs. The process was repeated 1,000 times to obtain 1,000 candidate sets. After that, predicted by FSM, the drug combinations with occurrences greater than 950 (*P*-value < 0.05) were chosen as the final drug combination datasets for a disease.

### 2.6 Performance evaluation

The area under the receiver operating characteristic curve (AUC) and the area under the precision-recall curve (AUPR), both widely used in various fields [24,25] were applied to evaluate our method. Given the severe class imbalance in the drug combination dataset, AUC and AUPR are not ideal metrics for evaluating classifiers on such datasets [26]. In this paper, in addition to AUC and AUPR, we also employed the Normalized Discounted Cumulative Gain (NDCG) as the primary evaluation metric. NDCG emphasizes the early retrieval performance of models and measures how closely a particular ranking resembles the ideal ranking [27]. Further information on this metric can be found in Supplementary Note 3. It’s important to note that, unless specified otherwise, the performance reported in this article is based on the average performance obtained from 100 rounds of 2-fold cross-validation.

## 3 Results

In this section, we first confirm that the “friendship feature” is an effective characteristic for representing drug information, applicable across multiple levels and heterogeneous data types. Subsequently, by analyzing the similarity network of friendship features among drugs, we identify specific patterns for single drugs that can be used to construct drug combinations. Using this property, we find that predicting these combinable drugs can reduce the search space for potential drug combinations. Following this, we propose a novel combination drug prediction model (FSM) and systematically compare its performance with several established models. Finally, we observe that the drug combinations predicted by our approach contain more effective combinations, fewer adverse effects, and maintain a moderate level of similarity within their friendship feature. Detailed results are provided below.

### 3.1 Friendship Feature: An Enhanced Representation and Integration Metric for Drug Characteristics

In recent years, various models have been developed under the assumption that “two drugs producing similar effects can exhibit synergy when used together [28].” These models aim to predict effective drug combinations based on drug similarity or distance. In our study, we departed from the conventional approach by considering the similarity of a drug’s relationships with all other drugs, as opposed to solely the relationship between the two drugs themselves (direct similarity).

To illustrate this concept, we used drug target data as an example. Separation distance is a common metric used to analyze the relationships among drug-target modules and disease modules within the Protein-Protein Interaction (PPI) network [11,29]. When two drugs, *a* and *b*, have a direct similarity (Separation distance) of *X*, their relationship can take on various forms (e.g., with drug a as the center and *X* as the radius), making it challenging to distinguish them from other drug combinations with the same distance (Fig. 1a, left). Consequently, representations of drug combinations based solely on direct similarity are vague and incomplete.

**Fig. 1.**
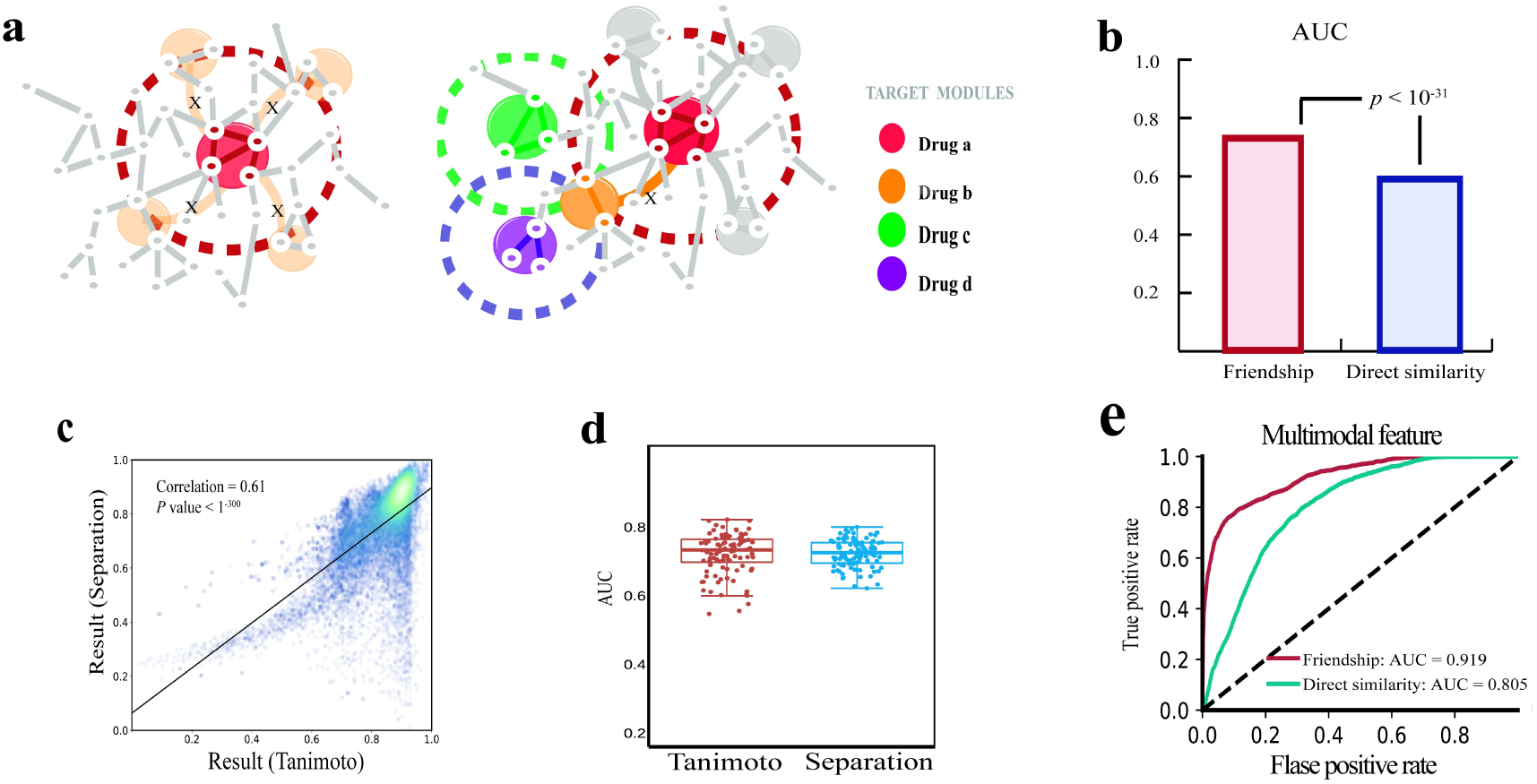
Friendship feature can accurately represent drug associations. (a) Schematic diagram of the advantage of friendship feature. (b) The performance comparison of Friendship feature and direct similarity on hypertension dataset. (c) Scatter plot of drug similarity based on separation distance and Tanimoto coefficient on hypertension dataset. (d) The performance comparison of different friendship features (separation distance and Tanimoto coefficient) on the hypertension dataset. (e) Comparison of the performance of friendship feature and direct similarity feature in multilevel data integration models.

However, we observed that integrating the relationship similarity between the two drugs and all other drugs could provide a more precise description of their relationship (Fig. 1a, right). Hence, we hypothesized that constructing higher-quality features by integrating interaction information related to a drug’s relationships with other drugs (Method) could be achieved, and we named this feature the “friendship feature.” To validate the efficacy of the friendship feature, we built a classifier using One-Class Support Vector Machines to predict drug combinations for cancer and hypertension, respectively. Simultaneously, we constructed a prediction model based on direct similarity (Separation distance). Our results indicated that the model based on the friendship feature significantly outperforms the one based on direct similarity (Separation distance) for both hypertension (Fig. 1b) and the cancer dataset (Supplementary Fig. 1). These findings support our hypothesis that using the relationships of each drug with all other drugs as features results in higher-quality and more distinctive drug descriptions, substantially improving predictive accuracy.

The above results confirm the effectiveness of the friendship feature when applied to drug target data based on the PPI network. However, the challenge lies in calculating the separation distance for other data types, limiting our ability to extend the use of the friendship feature to diverse data categories. To explore the adaptability of the friendship feature to other data types, we employed the Tanimoto coefficient as a similarity measure and utilized it to construct the friendship feature for drugs. Subsequently, we developed a drug combination prediction model based on this novel friendship feature. The scores of drug combinations generated by the two models, one using the friendship feature based on the separation distance and the other on the Tanimoto coefficient, displayed a strong overall correlation (Fig. 1c, Supplementary Fig. 2). Moreover, our results demonstrate that both friendship features exhibit highly consistent prediction capabilities (Fig. 1d, Supplementary Fig. 3), with the model employing the friendship feature based on the Tanimoto coefficient performing even better. Overall, these findings illustrate that the friendship feature of drugs, based on the Tanimoto coefficient, can capture similar information to the friendship feature based on the separation distance in the PPI network. This suggests the potential for extending the utility of the friendship feature to other data types.

As we know, various techniques can be used to characterize small molecules by examining genome, chemical structure, clinical, and pharmacological data. A recent study has revealed that similarity scores calculated for different data types exhibit little overall correlation, highlighting that each type of data measures different aspects of molecular activity [14]. Furthermore, integrating information from multiple sources has been shown to enhance the analytical power of models [13]. To assess whether the friendship feature effectively integrates data from different levels, we employed the friendship feature based on the separation distance of drug targets, pathway similarity, clinical similarity, chemical similarity, and adverse similarity to construct drug combination prediction models for cancer and hypertension datasets, respectively. Simultaneously, we utilized the friendship feature that integrates all similarity information to build the prediction model (Method). For comparison, the direct similarity feature was used to create corresponding single-feature models and integrated feature models. The results consistently demonstrate that, at each individual data level, the friendship feature outperforms direct similarity in predictive performance (Supplementary Table 1 and Supplementary Fig. 4a-d). Furthermore, whether integrating similarity information from different levels directly or employing the friendship feature to integrate information from various levels, both approaches outperform the use of a single feature (Supplementary Table 1 and Supplementary Fig. 4e), with the integrated model of the friendship feature delivering superior results in comparison to the direct similarity model (Fig. 1e).

### 3.2 Drugs That Can Form Combination Drugs Have Drug Combinability

Drug association analysis is a crucial step in understanding drug interactions and their impact on disease mechanisms [11,30]. We suggest that exploring drug-drug associations through the lens of “friendship similarity” can shed light on effective drug combinations and improve drug combination prediction. Utilizing the five data types mentioned earlier, we calculated a multimodal friendship feature for each drug. Pearson Correlation Coefficients (PCC) were employed to quantify the similarity between the 305 drugs. We constructed a drug-drug network by selecting drug pairs with correlations in the top 5% quantile, and assigned PCC values to the weighted edges. Community detection using the affinity propagation algorithm [31] revealed 25 communities, each represented by a specific color (Fig. 2a). Additional details about these communities can be found in Supplementary Table 2.

**Fig. 2.**
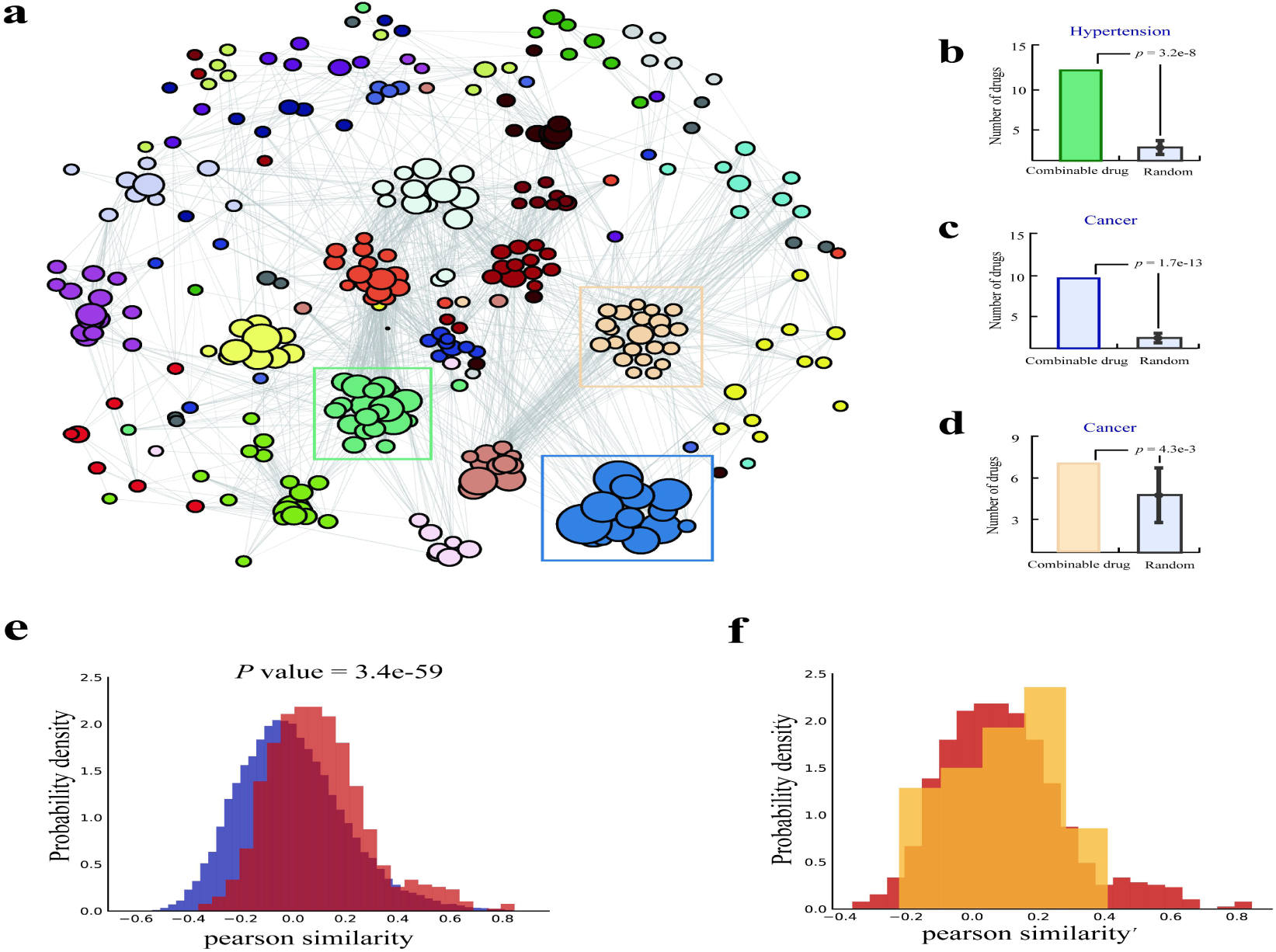
Drug-Drug Relationships Based on the Friendship Feature. (a) The network depicts drug interactions, with each color denoting a specific community. (b-d) Enrichment of combinable drugs for specific diseases (Cancer or Hypertension) within the specific community of the Drug-Drug Network. Gray boxes represent random expectations, and p-values are calculated using permutation tests (Supplementary Note 2). (e) Distribution of Pearson similarity between combinable drugs (in red) and other drug pairs (in blue) in the hypertension data set. (f) Distribution of Pearson similarity between effective drug combinations (in yellow) and combinable drugs (in red) in the hypertension data set.

Our observations indicate that drugs that form effective combinations for specific diseases, which are defined as combinable drugs, tend to cluster within the same community (Fig. 2a-d). PCC analysis revealed that combinable drugs exhibited significantly higher correlations compared to other drugs (Fig. 2e, Supplementary Fig. 5a), suggesting a preference for disease-specific combinable drugs to closely associate. Prior studies [11] have demonstrated that drugs or combinations whose target modules are in proximity to disease genes are more likely to be therapeutically effective. We also found that combinable drugs exhibit significant similarities. Disease constraints contribute to this phenomenon. At the drug target level, disease genes must be close to the combinable drug’s target module, resulting in the similarity of the drug’s friendship feature. We refer to this constraint as “drug combinability.”

We further analyzed whether the correlation between individual drugs within each drug combination is the highest among all possible combinations. Our findings revealed that effective drug combinations do not exhibit the highest Pearson similarity (Fig. 2f, Supplementary Fig. 5b), challenging the notion of drug pairs being infinitely close to each other. This observation aligns with the concept of “Complementary Exposure” in drug pairs, as described in a previous study [32].

### 3.3 Predicting Disease-Specific Combinable Drugs Can Facilitate Combination Drug Prediction

Given that the combinability of drugs can characterize whether a single drug is likely to be part of a combination therapy for a specific disease, predicting these single drugs, referred to as ’combinable drugs,’ in advance can reduce the search space for drug combinations and further enhance the efficiency of combination drug screening. We demonstrated the effectiveness of drug combinability using the OCSVM algorithm to predict combinable drugs for cancer and hypertension. Different performance indexes were employed (Fig. 3a-c, Supplementary Fig 6a-c), showing the algorithm’s ability to capture combinable drugs. We also assessed the ability of drug combinability to narrow down the search space, with results in Fig. 3d and Supplementary Fig. 6d. By using various thresholds to determine combinable drugs, we found that machine learning algorithms can predict disease-specific combinable drugs, significantly reducing the need for experimental pre-screening of existing drug pairs.

**Fig. 3.**
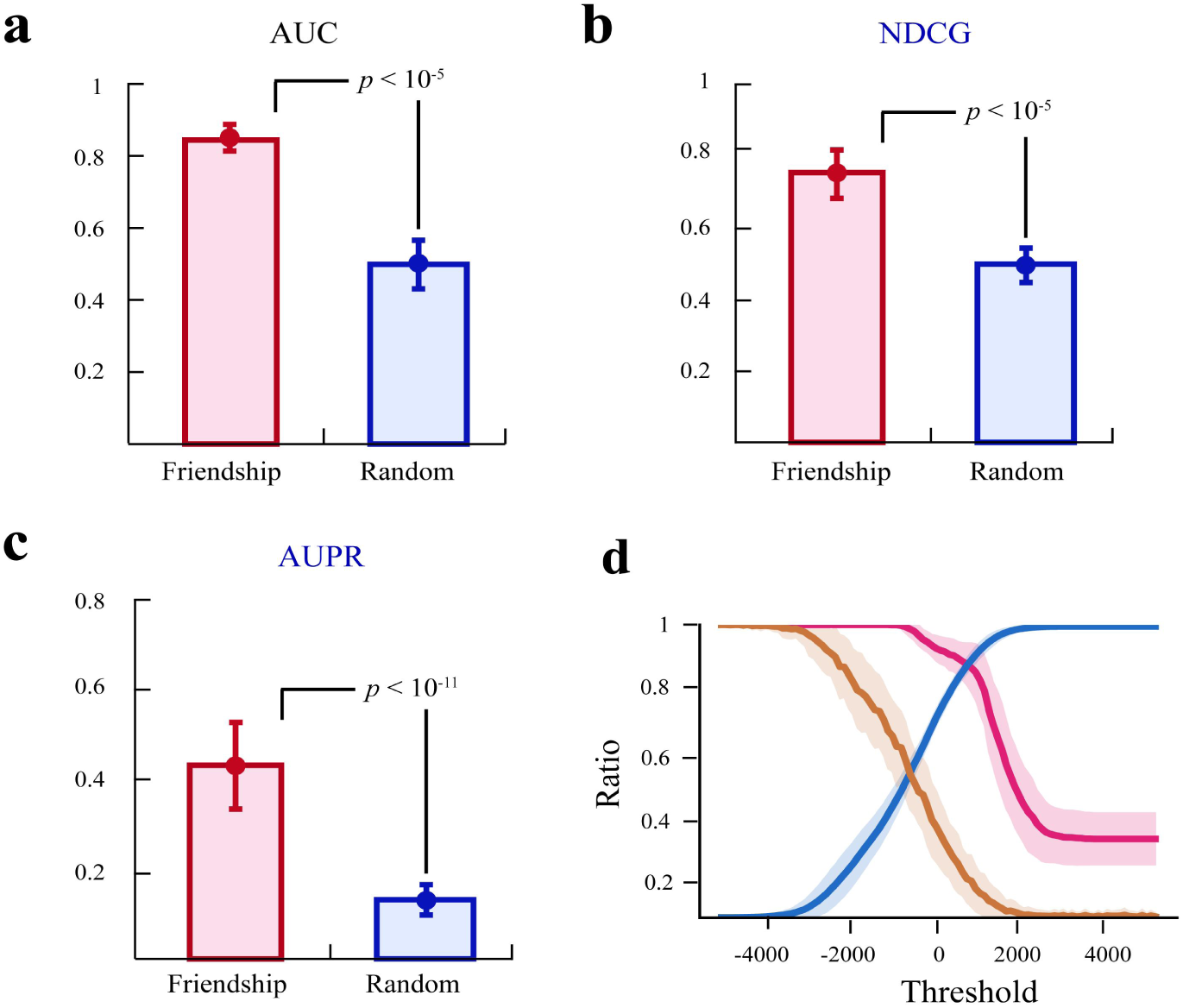
Demonstrating the Efficacy of Drug Combinability. (a-c) Highlighted in red are the predictive performances of combinable drugs for hypertension. The blue boxes (Random) depict random expectations (refer to Supplementary Note 2). (d) Illustrating the screening capacity of drug combinability within the drug space of the hypertension dataset. The blue line represents the ratio of the filtered space to the total space under various thresholds, and the yellow line shows the random expectation. The shaded region portrays the standard deviation based on 200 simulations.

### 3.4 The mainframe of FSM

Based on the findings above, we developed a new computational pipeline, called FSM, to identify efficacious combination therapies for a specific disease (Fig. 4). FSM integrates multiple types of drug data and interaction information between drugs by Friendship feature to obtain informative vector representations of drugs and drug combination. We then used OCSVM to predict the combinability of drugs, thereby filtering out the drug pairs that contain uncombinable single drugs. After that, FSM predicts the remaining drug pairs by using the OCSVM as classifier and friendship as feature. More details of the FSM pipeline can be found in Methods and supplementary Note 4.

**Fig. 4.**
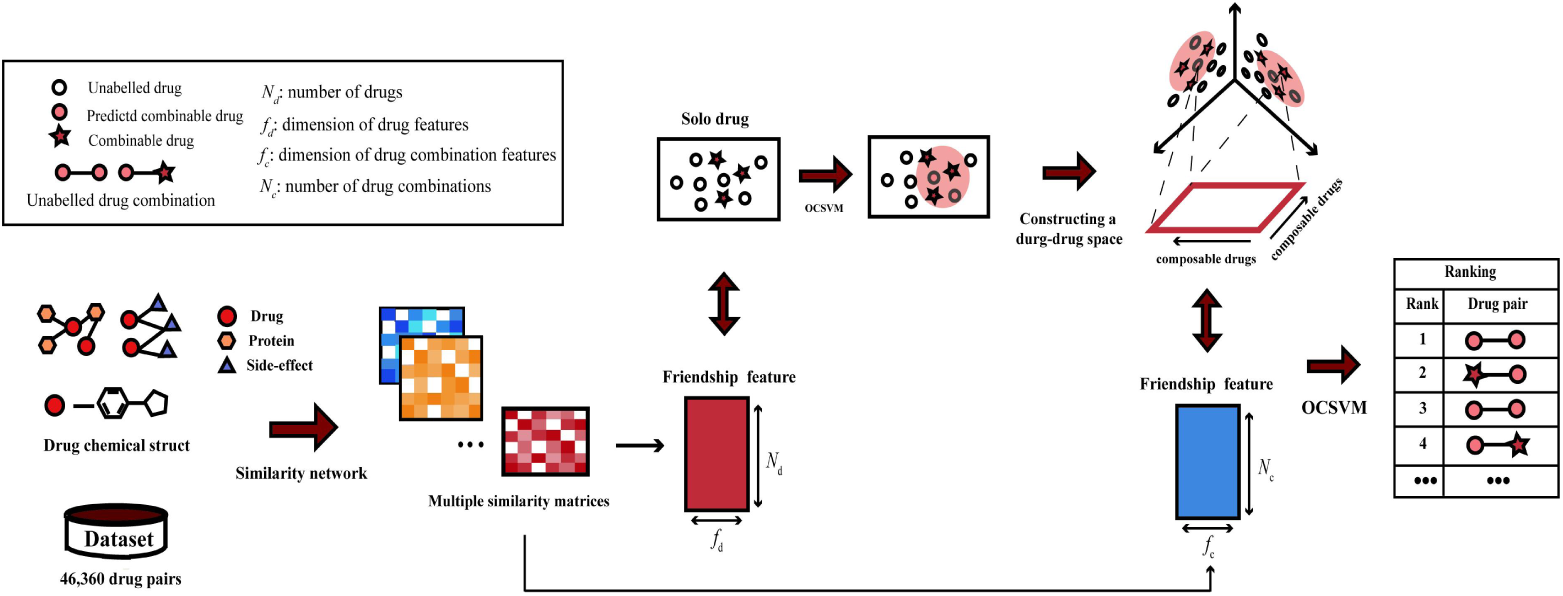
Flowchart of the FSM. Initially, FSM utilizes various drug-related data sources to compute multiple drug-drug similarities and incorporates interaction information between drugs to derive the friendship feature of both individual drugs and drug pairs. Subsequently, FSM employs the OCSVM algorithm to pinpoint combinable drugs specific to a given disease and establishes the combinable drug space to eliminate non-combinable drugs. Finally, FSM assesses the synergistic potential of the remaining drug pairs using the friendship feature and the OCSVM algorithm.

### 3.5 FSM Outperforms State-of-the-Art Methods

To evaluate the prediction power of FSM, we applied FSM on cancer and hypertension, two diseases with a large number of FDA-approved drug combinations, and compared FSM with five state-of-the-art methods, including NEMN [20], TLMCS [17], Li’s [18], Zhao [16], and Separation [11]. To obtain a reliable result, cross-validation and multiple evaluation metrics were used to evaluate the performance of the models. Our method displayed the best performance on the two datasets (Fig. 5a, b), indicating that FSM can guarantee excellent performance while directly filtering most of the search space. This also proves that exploring the combinability of drugs is a promising route to overcome combinatorial explosion. In addition, compared with TLMCS that uses original drug features or other methods based on direct similarity, the AUC, AUPR, and NDCC of FSM are higher, which proved that the friendship feature can better characterize the relationship of drug-drug and notably improve the ranking results.

**Fig. 5.**
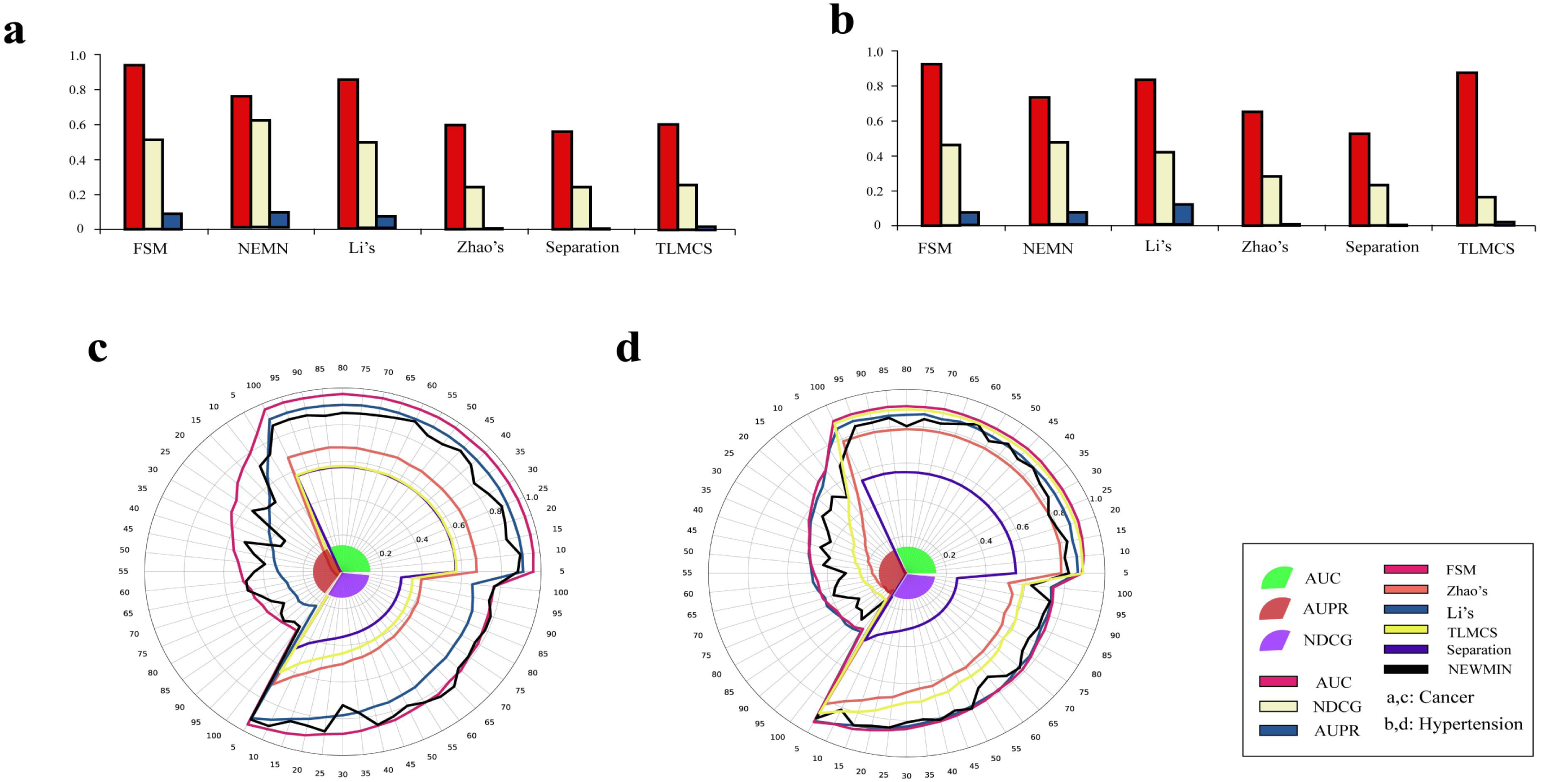
Performance Comparison of Prediction Models on Hypertension and Cancer Datasets. Subfigure (a) shows the AUPR, AUC, and NDCG values for the five methods on the cancer dataset, while subfigure (b) displays the corresponding results for the hypertension dataset. Subfigures (c) and (d) present performance comparisons between FSM and other state-of-the-art methods on imbalanced datasets (The labels on the rings represent the negative-to-positive sample ratio) for the cancer dataset and the hypertension dataset, respectively.

As we know, the ratio of the positive samples and negative samples can affect the performance of machine learning algorithms [33]. In the meanwhile, in real datasets for drug combination prediction, the ratio is highly imbalanced. To further validate our method, we randomly extract the negative sample from the original dataset to construct a new sample space, with the ratio of positive and negative samples from 1:5 to 1:100 (step = 5). The results (Fig. 5c, d) showed that the performance of all methods decreased with the increase of negative samples. In the meanwhile, our model still has a higher performance than other methods. Thus, the noticeable performance improvement over other prediction methods demonstrated the superiority of FSM.

### 3.6 Drug Combinations Predicted by FSM Exhibit Moderate Friendship Similarity and Lower Adverse Effects

By FSM, 879 drug combinations were identified for cancer and 499 drug pairs were obtained for hypertension (Supplementary Table 3 - 4). As described above (Fig. 2f, Supplementary Fig. 5b), the drug pairs of clinical drug combinations have moderate similarities. Therefore, we also investigated whether the predicted drug combinations obeyed the rule. The Pearson similarities of the drug pair predicted by FSM, the drug pairs predicted as negative by FSM, and the drug combinations composed of combinable drugs predicted by FSM were calculated respectively. In both hypertension (Fig. 6a) and cancer datasets (Supplementary Fig. 7a), the PCCs of the predicted drug combinations are significantly higher than the negative drug pairs (Kolmogorov-Smirnov test, *P*-value = 1.2e-315). In the meanwhile, the drug combinations predicted by FSM do not contain the ones with particularly large Pearson similarity (Fig. 6b, Supplementary Fig. 7b). These results indicate that our predicted drug combinations follow the pattern of the effective drug combinations identified above (moderate similarities).

**Fig. 6.**
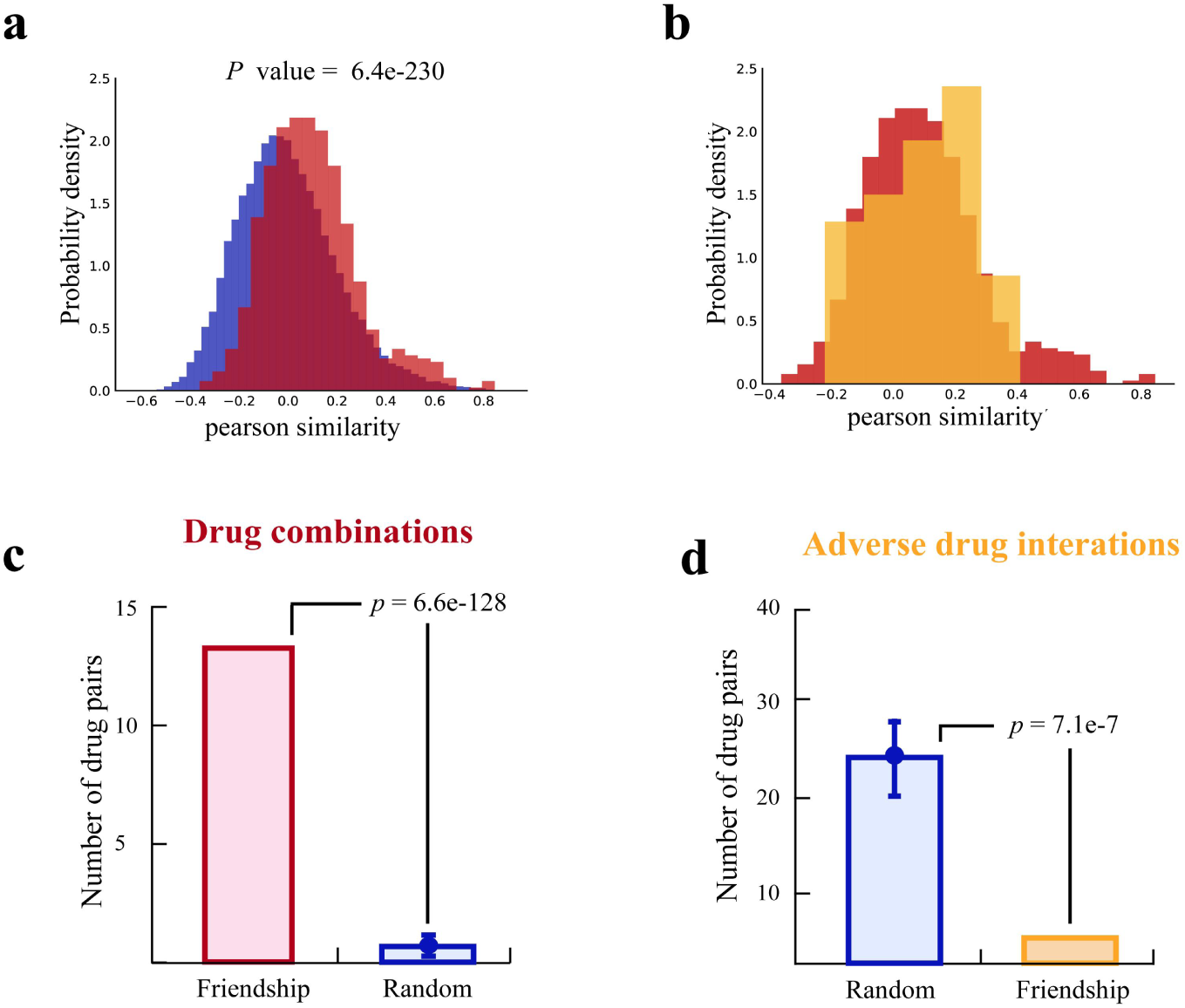
Combination drugs predicted by FSM in the hypertension dataset. (a) Distribution of Pearson similarities between combinable drugs (in red) and similarities of other drug pairs (in blue). (b) Distribution of Pearson similarity between drug combinations (in yellow) predicted by FSM and similarities of all combinable drugs (in red). (c, d) Enrichment results of the predicted drug combinations with known drug combinations (c) and drug combinations with side effects (d). The color histogram represents the predicted drug combinations (in pink) and clinically reported adverse drug interactions (in yellow). The blue box indicates the intersection with an equal number of randomly selected drugs.

In addition, we also investigated whether the predicted drug combinations could be enriched with more effective drug pairs and less adverse drug pairs. On hypertension data set, comparing with random drug pairs (Supplementary Note 2), the drug combinations predicted by FSM have more overlaps with gold-standard drug pairs (*P*-value = 6.6e-128, Fig. 6c) and fewer overlaps with adverse drug pairs (*P*-value = 7.1e-7, Fig. 6d), and the same conclusion could be drawn on cancer data set (Supplementary Fig. 8a and Supplementary Fig. 8b). All these results indicate that FSM could facilitate the screen of effective drug pairs, and would help reduce adverse due to drug-drug interactions. It is consistent with the principle of non-overlapping pharmacology in drug combination design [32].

All these results demonstrated that FSM follows the extended combination drug design principle, which can significantly enrich effective drug combinations and could avoid toxicity caused by drug interactions. Furthermore, FSM has successfully identified drug combinations that have been experimentally verified and can guide future experimental validation and prospective clinical trials.

## 4. Discussion

Combinational therapy has been widely explored as a major route to overcome complex diseases [1]. However, the current methods to identify combination drugs for specific diseases still have some limitations. To solve these problems, an effective model, denoted as FSM, was proposed in this study. Through comprehensive tests on hypertension and cancer datasets, it is proved that FSM can achieve markedly performance improvement over other state-of-the-art prediction methods. FSM is based on the Friendship feature, which can represent the drugs and drugs pairs more precisely. Furthermore, our model is implemented by OCSVM, which overcomes the intrinsic bias of the former approaches. Notably, we found that effective drug combinations exhibit an interaction pattern in high-dimensional, heterogeneous feature space (Moderate similarity), and these findings provide new insights for the discovery of drug combinations.

Network-based approaches have already offered a promising framework to identify novel insights to accelerate drug discovery [34]. Previous studies have discussed the molecular pattern underlying the targets of drugs on networks and applied it to predict effective drug combinations. However, we discovered the previous methods are not good enough to provide enough information to accurately describe the drug combination in the face of a vast combinatorial space. In this manuscript, we proposed a friendship feature and confirmed that the feature offered a powerful representation of drug-drug relationships. Through drug-drug association analysis, we propose that the disease has a restriction on its combinable drugs, and prove that it can be well captured by machine learning algorithms. These results verify that we can filter most of the search space in advance by removing the uncombinable drugs, thereby alleviating the problem of drug combination explosion.

In the present study, although we proved that the Friendship feature can be extended to different data types, and FSM was established by multiple data sources. However, there is still a lot of data that could be incorporated to increase the performance of the computational model. For example, Sun et al. [8] found that the gene expression data may reflect the underlying mechanism of the synergistic effect. Therefore, with the acquisition of new data, our model would describe drugs more comprehensively and be more accurate for predicting drug combinations for specific diseases. Furthermore, with the increase in the types of drugs, the dimension of the friendship feature will also expand. How to use feature engineering to obtain low-dimensional but informative representations of drugs remains the goal of our future work.

In summary, we show herein the power of Friendship features and formulated an efficient model (FSM) for the prediction of clinically efficacious drug combinations for specific diseases. In addition, the model cuts in from the perspective of single-drug combinability, allows efficient large-scale search space screening, to accelerates the identification of combinational therapy.

## Code and data availability

The code and data are available at https://github.com/nbnbhwyy/FSM.

## Acknowledgments.

This work was supported by the Fundamental Research Funds for the Central Universities to X. Z. (2662023XXPY003).

## SUPPLEMENTARY INFORMATION

### Supplementary Note 1: Data sets

drug targets: The drug-target data was extracted from Cheng et al. [1]. They obtained comprehensive drug target linkages by integrating high-quality physical drug-target interactions of FDA-approved and clinically investigational drugs from six commonly used data sources, which contains 15,051 drug-target interactions connecting 4,428 drugs and 2,256 human targets.

Adverse effects: Side effects were downloaded from the OFFSIDES database [2], which includes 438,801 associations connecting 1,332 drugs and 10,097 adverse events.

Chemical structures: All the chemical structure information (SMILES format) was downloaded from the DrugBank database [3].

Pathway: The Pathway data was downloaded from the website: http://www.gsea-msigdb.org/gsea/index.jsp. The pathway data, including 186 Human functional pathways, covers 5,266 human genes.

Clinical data: The drug clinical information was represented by the code of the Anatomical Therapeutic Chemistry (ATC) classification system and all drugs were collected from the DrugBank database [3].

Gold standard data: The gold standard drug combinations were derived from the work of Cheng et al. [1], which includes 681 unique pairwise drug combinations composed of 362 drugs. Our study selected the possible combinations among all drugs of the golden standard for pairwise drug combinations as a search scope of the model, including 362 drugs and 65,341 pairwise drug combinations in total. After that, only the drugs with pathways, adverse effects, targets, and structural information were kept. Finally, we constructed a data set of 305 drugs and 46,360 pairwise drug combinations. To verify the prediction function of the model, we focus on hypertension and cancer, two diseases with many FDA-approved pairwise combinations.

Adverse drug-drug interactions: The adverse data was collected from the DrugBank database [3]. We extracted adverse reactions caused by drug combinations in our dataset. In total, 3,198 clinically-reported adverse connecting 254 unique drugs were retained.

### Supplementary Note 2: Permutation tests

The permutation test is a standard test to compute statistical significance [4]. The null hypothesis of the permutation test is generated by counting all possible values of the test statistic under rearrangements of the true labels on the shuffled data points. Previous studies [5] have found that thousands of test statistics from the permutation distribution usually are sufficient to get an accurate estimate of the exact P-value. Specifically, if M test statistics, ti, i = 1, …, M, are randomly sampled from the permutation distribution, a one-sided Monte Carlo P-value for a test that rejects for large values of t is p, and the statistical significance is given by:

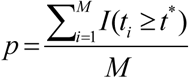

Where *t\** is an observed value. A nominal P is computed by counting the number of shuffled differences greater than or equal to the true difference.

For the P-value in Fig. 2a-e, we randomly shuffled the gold standard (Combinable drugs) and repeated this randomization process 100 times. The P-value is obtained by calculating the possibility of the random value no less than the true value.

For the P-value in Fig. 3b-d, if the number of intersection drugs between combinable drugs and a community is *X.* we randomly selected the same number of drugs as combinable drugs from all possible drugs and we performed 1,000 randomizations. For these 1,000 randomizations, the possibility of the overlaps between the random drugs and the communities no less than X was set as the p-value. We performed the same permutation test for the construction of the pathway feature.

For the P-value in Fig. 4a-c, we randomly shuffled the gold standard (Combinable drugs) and repeated this randomization process 1000 times. The P-value is computed by calculating the possibility of random value no less than the true value.

For the P-value in Fig. 7c,d, we randomly shuffled the gold standard (Combinable drugs) and repeated this randomization process 1000 times. The P-value is computed by calculating the possibility of random value no less than the true value.

### Supplementary Note 3: Normalized Discounted Cumulative Gain (NDCG)

We use NDCG metric [6] to compare the quality of the ranks given by each method. This score shows how close a particular rank is to the ideal rank. In this work, the ideal ranking is the one where all the FDA-approved drug combinations for a disease are on the top of the ranked list. Suppose the total number of drug pairs is m, the metric NDCG for a particular ranked list is defined as:

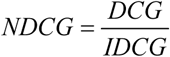

In this equation, DCG represents the Discounted Cumulative Gain of the ranking we want to score, while IDCG represents the discounted cumulative gain of the ideal ranking. The value of NDCG is between 0 to 1, where a value of 1 represents the case in which the cumulative gain of the given ranking is equal to the cumulative gain of the ideal ranking. The DCG for a given ranking can be calculated as follows:

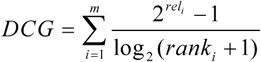

In equation 3, rel is the relevance value, which is defined as a binary value: 1 for drugs FDA-approved for the given disease, and 0 for all other drugs).

### Supplementary Note 4: One-class SVM

One-class SVM (OCSVM) [7], the support vector model is trained on data that has only one class. This method infers the properties of known instances from these properties and can predict where unknown instances are like or unlike the known examples. OCSVM is especially appropriate for the predicting task, which gives a few positive instances and many unlabeled instances.

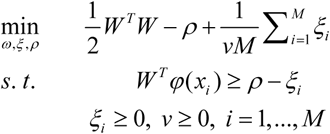

where φ(x_i_) maps xi into a higher-dimensional space. Due to the possible high dimensionality of the vector variable w, we usually turn the above problem to its dual problem to solve.

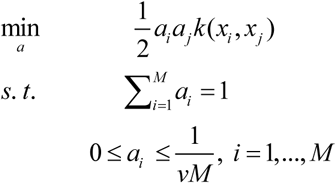

In this work, we use OCSVM as the classifier, and select the linear function and radial basis function as the kernel function for predicting drug composability and potential drug combination, respectively.

### Supplementary Figures

**Supplementary Fig. 1.**
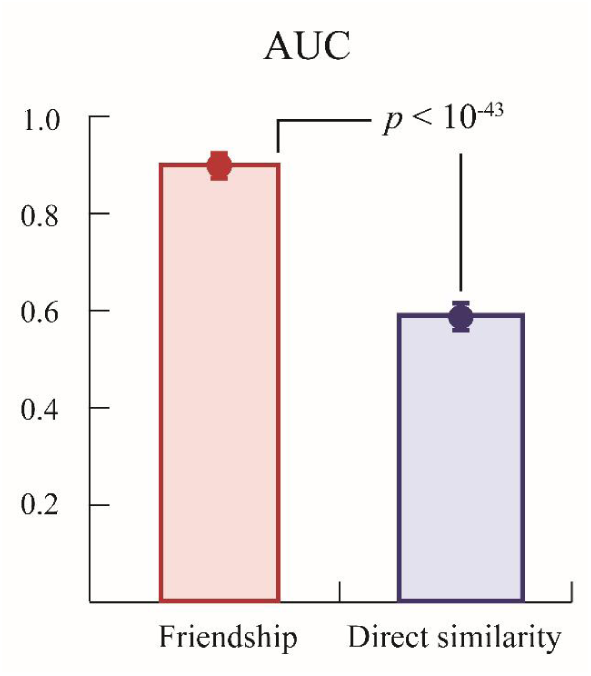
The performance (AUC) comparison of Friendship feature and direct similarity on cancer dataset.

**Supplementary Fig. 2.**
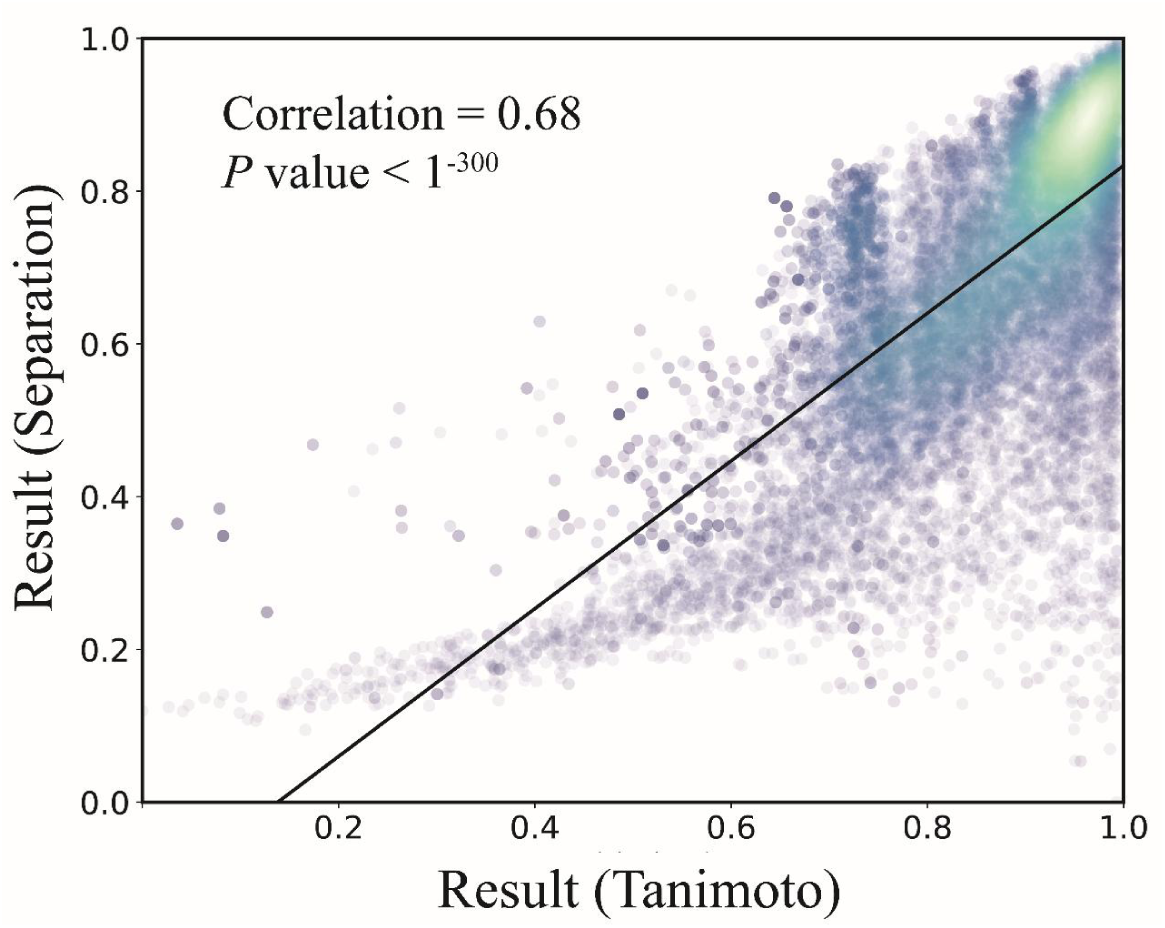
Scatter plot of drug similarity based on separation distance and Tanimoto coefficient on cancer dataset.

**Supplementary Fig. 3.**
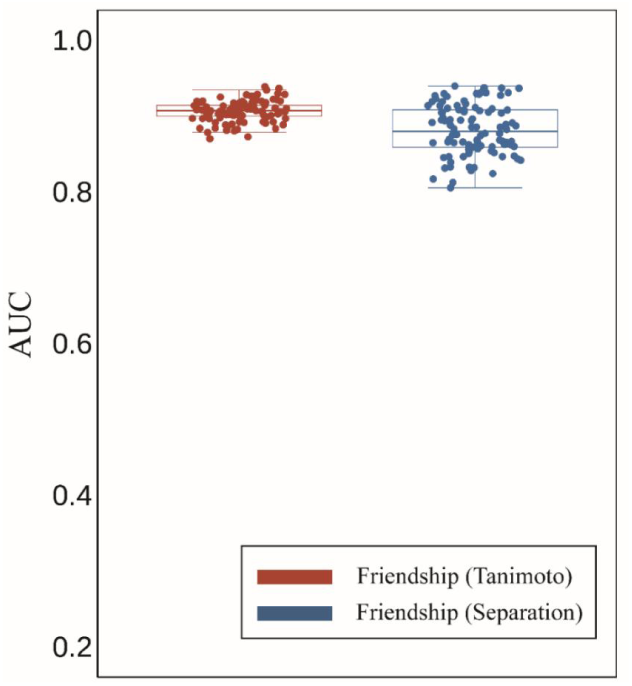
The performance comparison of different friendship similarities (separation distance and Tanimoto coefficient) on cancer dataset.

**Supplementary Fig. 4.**
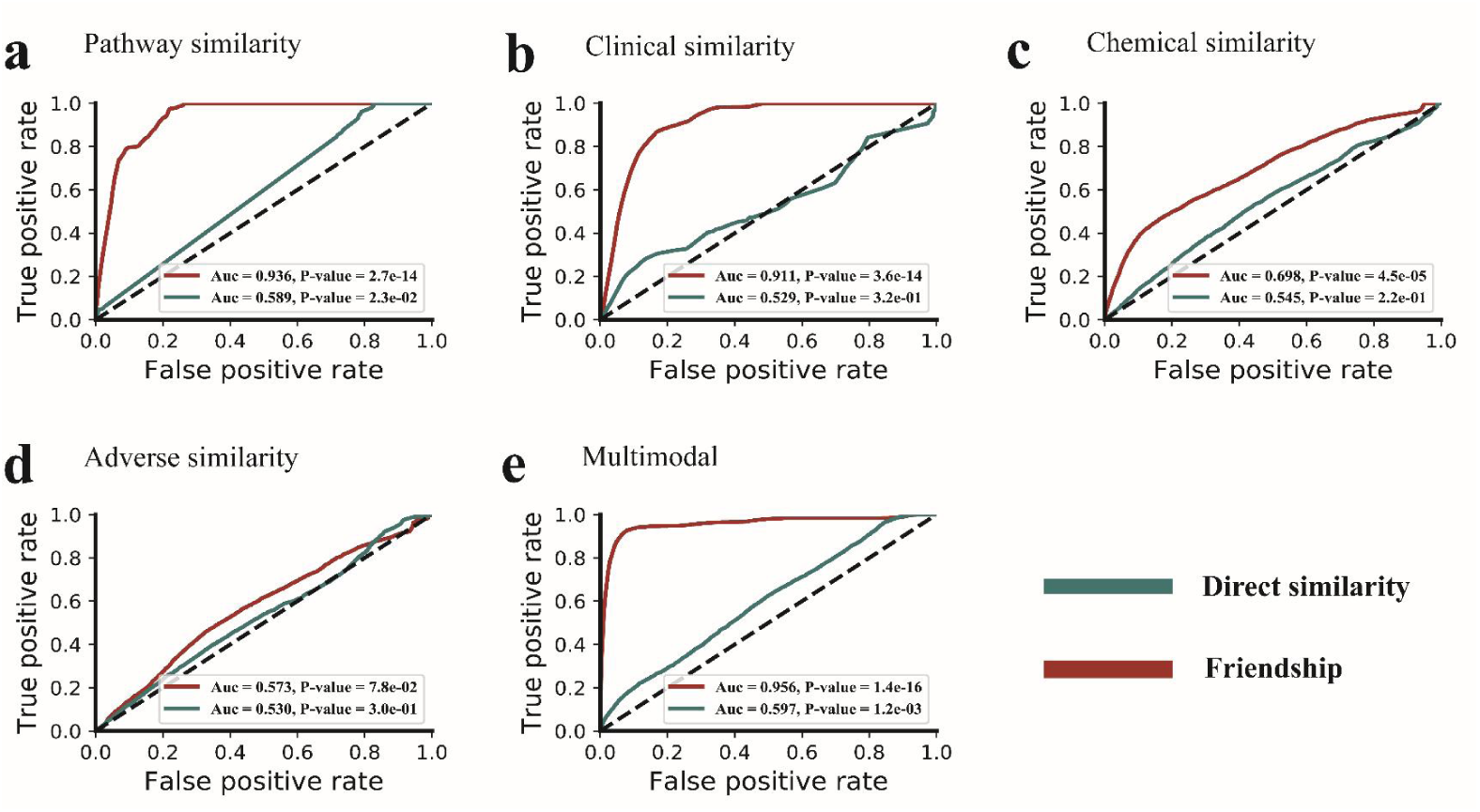
Performance comparison of friendship feature and direct similarity on the cancer dataset (different data types).

**Supplementary Fig. 5.**
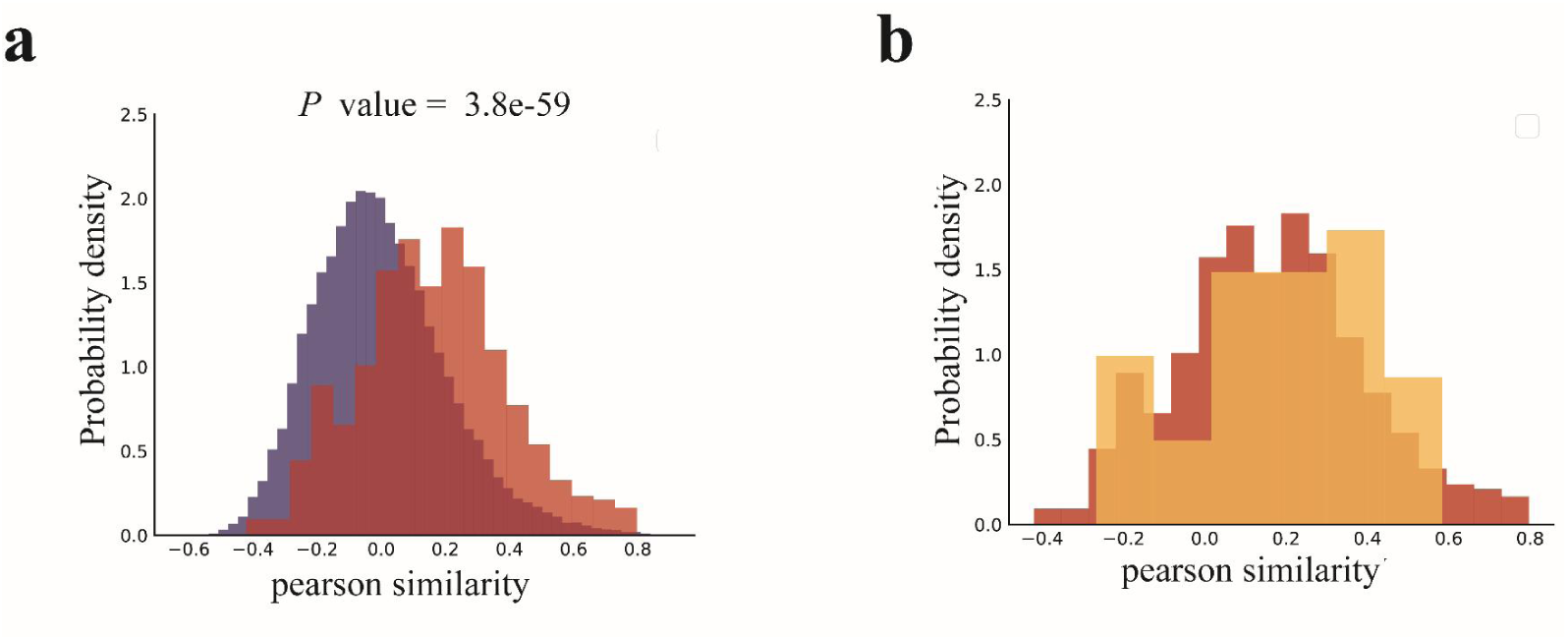
The similarity distributions of effective drug combinations on the cancer dataset. (a) The distribution of Pearson similarities between the drugs of combinable drugs (red) and Pearson similarities between other drug pairs (blue). (b) The distribution of Pearson similarities of effective drug combinations (yellow) and combinable drugs (red).

**Supplementary Fig. 6.**
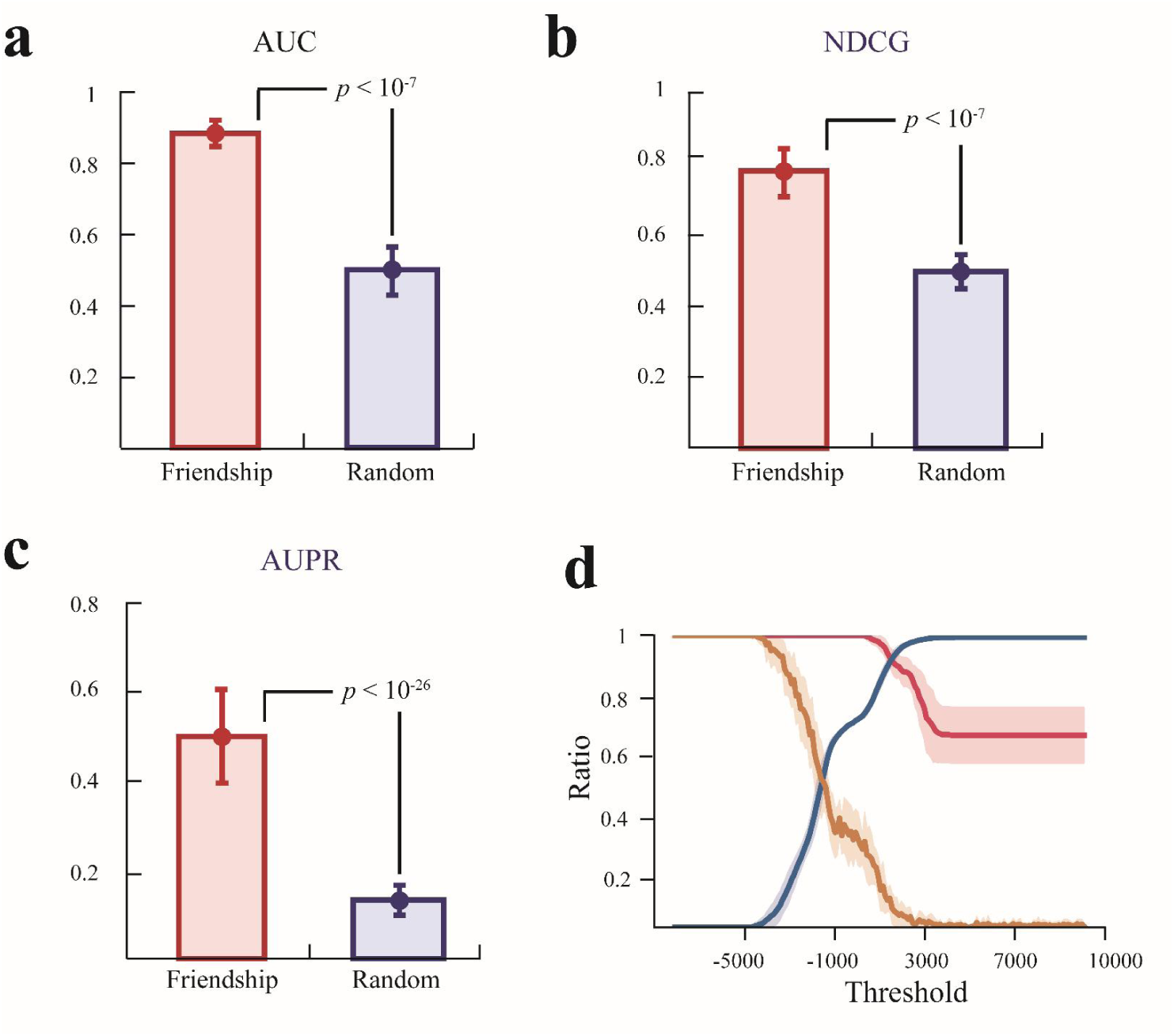
The utility of the composability of drugs. (a-c) The red boxes show the predictive performance of cancer combinable drugs. Blue boxes (Random) show the random expectations. (d) The screening ability of drug composability on the cancer dataset. The blue line represents the ratio of the filtered space to the total space under different thresholds; the red line indicates the ratio of the remaining drug combinations to all drug combinations under different thresholds; yellow lines show the random expectation, and the transparent range represents the standard deviation of 200 simulations.

**Supplementary Fig. 7.**
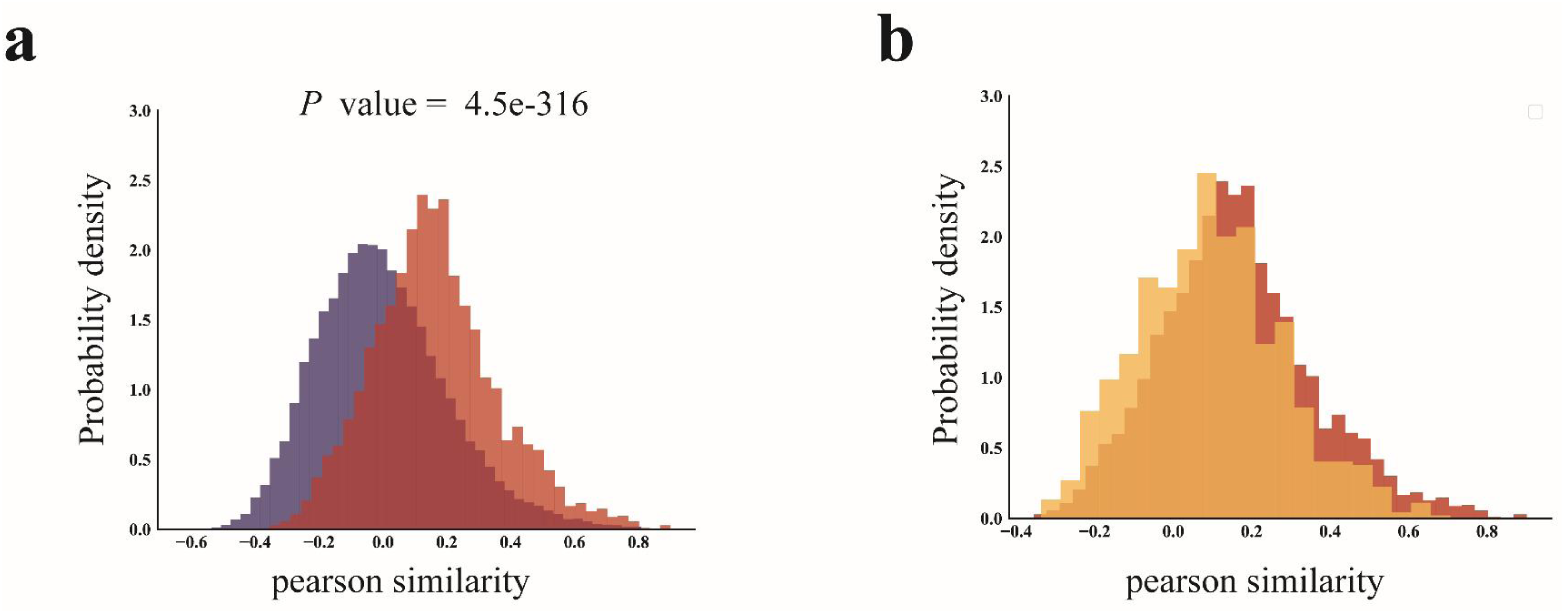
The similarity distributions of predicted drug combinations on the cancer dataset. (a) The distribution of Pearson similarities between the combinable drugs (red) and Pearson similarities between the other drug pairs (blue) on the cancer data. (b) The distribution of Pearson similarities between effective drug combinations (yellow) and Pearson similarities between the combinable drugs (red) on the cancer data.

**Supplementary Fig. 8.**
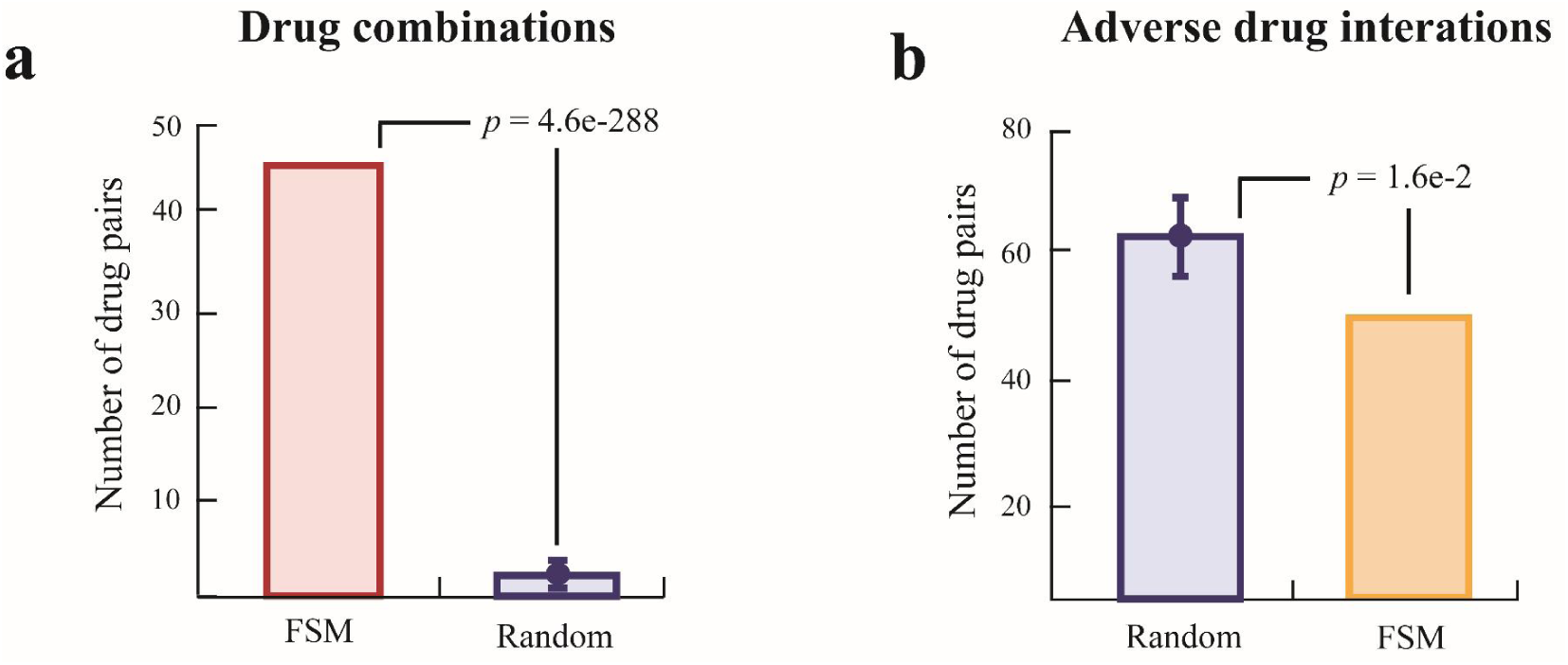
The combination drugs predicted by FSM on the cancer dataset. Enrichment results of the predicted drug combination with known combination drug (a), and drug combination with side effects (b). The color histogram shows the effective drug combination (pink) and the clinically reported adverse drug interactions (yellow). The blue box shows the intersection between the clinically reported adverse drug interactions and the same number of random drugs (with the predicted combinations by FSM).

### Supplementary Tables

**Supplementary Table 1.** Performance comparison of various feature.

|  | Pathway similarity |  | Clinical similarity |  | Chemical similarity |  | Adverse similarity |  | Multimodal feature |  |
| --- | --- | --- | --- | --- | --- | --- | --- | --- | --- | --- |
|  | AUC | P-value | AUC | P-value | AUC | P-value | AUC | P-value | AUC | P-value |
| Friendship similarity | 0.670 | 5.4e-03 | 0.858 | 4.4e-07 | 0.661 | 2.7e-03 | 0.841 | 7.9e-07 | 0.919 | 2.4e-09 |
| Direct similarity | 0.591 | 4.1e-02 | 0.531 | 3.5e-01 | 0.544 | 2.7e-01 | 0.528 | 3.2e-01 | 0.805 | 2.6e-03 |

**Supplementary Table 2.** The relationship between the drug module and combinable drugs.

|  | Number of drugs | Number of combinable drugs (Hypertension) | P-value | Number of combinable drugs (Cancer) | P-value |
| --- | --- | --- | --- | --- | --- |
| Cluster_1 | 7 | 3 | 1.7e-3 | 1 | 0.43 |
| Cluster_2 | 23 | 1 | 0.88 | 0 | 0.97 |
| Cluster_3 | 17 | 2 | 0.59 | 2 | 0.49 |
| Cluster_4 | 13 | 0 | 0.93 | 5 | 3.3e-3 |
| Cluster_5 | 9 | 0 | 0.87 | 0 | 0.87 |
| Cluster_6 | 18 | 2 | 0.58 | 0 | 0.95 |
| Cluster_7 | 9 | 2 | 0.17 | 1 | 0.49 |
| Cluster_8 | 13 | 0 | 0.95 | 0 | 0.91 |
| Cluster_9 | 9 | 0 | 0.90 | 0 | 0.86 |
| Cluster_10 | 8 | 1 | 0.49 | 0 | 0.87 |
| Cluster_11 | 9 | 0 | 0.86 | 0 | 0.87 |
| Cluster_12 | 9 | 1 | 0.61 | 1 | 0.54 |
| Cluster_13 | 14 | 1 | 0.73 | 10 | 1.7e-13 |
| Cluster_14 | 7 | 1 | 0.43 | 0 | 0.85 |
| Cluster_15 | 10 | 3 | 5.9e-2 | 0 | 0.88 |
| Cluster_16 | 21 | 0 | 0.97 | 0 | 0.94 |
| Cluster_17 | 9 | 1 | 0.58 | 1 | 0.51 |
| Cluster_18 | 12 | 0 | 0.91 | 1 | 0.58 |
| Cluster_19 | 11 | 3 | 6.1e-2 | 3 | 5.1e-2 |
| Cluster_20 | 14 | 2 | 0.49 | 2 | 0.42 |
| Cluster_21 | 16 | 9 | 3.2e-8 | 0 | 0.95 |
| Cluster_22 | 9 | 1 | 0.55 | 1 | 0.56 |
| Cluster_23 | 6 | 3 | 1.7e-3 | 1 | 0.42 |
| Cluster_24 | 23 | 3 | 0.48 | 7 | 4.3e-3 |
| Cluster_25 | 9 | 0 | 0.86 | 0 | 0.84 |

**Supplementary Table 3.** Drug combinations predicted by FSM (cancer data set)

| Drug A | Drug B | Score |
| --- | --- | --- |
| Bortezomib | Sirolimus | 212.7626 |
| Lapatinib | Ciclosporin | 197.1234 |
| Bortezomib | Bosutinib | 192.8361 |
| Bortezomib | Ciclosporin | 188.8564 |
| Pazopanib | Sirolimus | 186.8361 |
| Lapatinib | Sirolimus | 186.3324 |
| Vorinostat | Ciclosporin | 185.8876 |
| Bortezomib | Epirubicin | 183.2957 |
| Simvastatin | Lapatinib | 182.4397 |
| Sorafenib | Sirolimus | 179.8125 |
| Vorinostat | Diltiazem | 178.3768 |
| Vorinostat | Sirolimus | 177.5122 |
| Bortezomib | Afatinib | 176.7038 |
| Vemurafenib | Ciclosporin | 176.086 |
| Lapatinib | Hydrocortisone | 175.8059 |
| Pentamidine | Ciclosporin | 175.7239 |
| Pazopanib | Ciclosporin | 175.4676 |
| Bosutinib | Sirolimus | 174.7806 |
| Vemurafenib | Docetaxel | 174.7341 |
| Lapatinib | Docetaxel | 173.4558 |
| Lapatinib | Etoposide | 173.0814 |
| Nilotinib | Sirolimus | 172.5697 |
| Epirubicin | Imatinib | 172.1449 |
| Bortezomib | Etoposide | 171.7725 |
| Bortezomib | Hydrocortisone | 171.6215 |
| Sunitinib | Etoposide | 171.55 |
| Vemurafenib | Sirolimus | 171.4029 |
| Bortezomib | Topotecan | 171.0097 |
| Simvastatin | Dasatinib | 170.9769 |
| Simvastatin | Bortezomib | 170.8656 |
| Sirolimus | Pentamidine | 170.3887 |
| Sunitinib | Pentamidine | 169.0045 |
| Daunorubicin | Ciclosporin | 168.964 |
| Ciclosporin | Gefitinib | 168.8234 |
| Bortezomib | Vemurafenib | 168.3196 |
| Bortezomib | Docetaxel | 168.0676 |
| Bosutinib | Docetaxel | 167.9565 |
| Doxorubicin | Pazopanib | 167.3909 |
| Dasatinib | Etoposide | 166.4738 |
| Sunitinib | Sirolimus | 166.1875 |
| Sunitinib | Docetaxel | 166.1619 |
| Bosutinib | Hydrocortisone | 166.0836 |
| Bortezomib | Lapatinib | 165.9323 |
| Bosutinib | Ciclosporin | 165.5935 |
| Lapatinib | Warfarin | 165.2672 |
| Doxorubicin | Vemurafenib | 164.7256 |
| Topotecan | Sunitinib | 164.6332 |
| Vorinostat | Sunitinib | 164.0017 |
| Sorafenib | Ciclosporin | 163.4592 |
| Dasatinib | Ciclosporin | 163.2396 |
| Sirolimus | Erlotinib | 162.8192 |
| Lapatinib | Pentamidine | 162.6179 |
| Bortezomib | Vandetanib | 162.2976 |
| Sunitinib | Epirubicin | 162.2094 |
| Vemurafenib | Daunorubicin | 162.1045 |
| Lapatinib | Paclitaxel | 161.9897 |
| Bortezomib | Paclitaxel | 161.8218 |
| Docetaxel | Imatinib | 161.4079 |
| Bosutinib | Etoposide | 161.4009 |
| Vemurafenib | Etoposide | 161.1792 |
| Lapatinib | Vorinostat | 160.9329 |
| Hydrocortisone | Dasatinib | 160.6852 |
| Erlotinib | Ciclosporin | 160.463 |
| Sirolimus | Afatinib | 160.186 |
| Vorinostat | Erlotinib | 160.1826 |
| Doxorubicin | Sirolimus | 160.0623 |
| Sirolimus | Dasatinib | 160.0259 |
| Fludarabine | Sirolimus | 160.0197 |
| Vemurafenib | Paclitaxel | 159.7823 |
| Vemurafenib | Epirubicin | 159.6614 |
| Simvastatin | Pazopanib | 159.6542 |
| Vorinostat | Levofloxacin | 159.5848 |
| Vemurafenib | Erlotinib | 159.5686 |
| Bortezomib | Gefitinib | 159.5135 |
| Lapatinib | Epirubicin | 159.4705 |
| Daunorubicin | Sirolimus | 159.4648 |
| Bosutinib | Ezetimibe | 159.3714 |
| Bortezomib | Daunorubicin | 158.8732 |
| Lapatinib | Doxorubicin | 158.6934 |
| Sorafenib | Topotecan | 158.4102 |
| Bortezomib | Vinblastine | 158.0243 |
| Doxorubicin | Ciclosporin | 157.7908 |
| Lapatinib | Ezetimibe | 157.6992 |
| Celecoxib | Epirubicin | 157.3263 |
| Pazopanib | Epirubicin | 157.3244 |
| Fludarabine | Docetaxel | 157.1934 |
| Sirolimus | Imatinib | 157.1381 |
| Bortezomib | Diltiazem | 157.124 |
| Bortezomib | Quinidine | 157.0805 |
| Hydrocortisone | Afatinib | 156.8084 |
| Sunitinib | Ciclosporin | 156.6794 |
| Sorafenib | Vinblastine | 156.6563 |
| Paclitaxel | Gefitinib | 156.6112 |
| Doxorubicin | Vorinostat | 156.5298 |
| Simvastatin | Sunitinib | 156.4862 |
| Afatinib | Ciclosporin | 156.3432 |
| Docetaxel | Dasatinib | 156.2904 |
| Dasatinib | Warfarin | 156.235 |
| Celecoxib | Sirolimus | 156.0692 |
| Bosutinib | Ceftriaxone | 155.6989 |
| Lapatinib | Topotecan | 155.6481 |
| Fludarabine | Ciclosporin | 155.5873 |
| Bortezomib | Doxorubicin | 155.5313 |
| Lapatinib | Prednisolone | 155.4876 |
| Daunorubicin | Pazopanib | 155.0547 |
| Docetaxel | Afatinib | 154.847 |
| Nilotinib | Ciclosporin | 154.7481 |
| Vorinostat | Imatinib | 154.6476 |
| Topotecan | Erlotinib | 154.4868 |
| Lenalidomide | Pentamidine | 154.3993 |
| Sirolimus | Gefitinib | 153.9314 |
| Paclitaxel | Dasatinib | 153.8511 |
| Bosutinib | Rosuvastatin | 153.8133 |
| Lenalidomide | Daunorubicin | 153.1602 |
| Bosutinib | Daunorubicin | 152.7948 |
| Pazopanib | Hydrocortisone | 152.2637 |
| Simvastatin | Vemurafenib | 152.01 |
| Vemurafenib | Montelukast | 152.0043 |
| Etoposide | Gefitinib | 151.9804 |
| Sirolimus | Vandetanib | 151.9285 |
| Sirolimus | Gemcitabine | 151.8702 |
| Afatinib | Etoposide | 151.6855 |
| Bortezomib | Prednisolone | 151.678 |
| Sorafenib | Epirubicin | 151.6541 |
| Bortezomib | Ofloxacin | 151.4573 |
| Simvastatin | Sorafenib | 151.4021 |
| Daunorubicin | Docetaxel | 151.322 |
| Daunorubicin | Rosuvastatin | 151.2103 |
| Doxorubicin | Topotecan | 150.8362 |
| Bortezomib | Tirofiban | 150.4962 |
| Bosutinib | Paclitaxel | 150.3558 |
| Vemurafenib | Hydrocortisone | 150.2588 |
| Vemurafenib | Sunitinib | 150.1258 |
| Sunitinib | Hydrocortisone | 150.1083 |
| Hydrocortisone | Imatinib | 150.0235 |
| Nilotinib | Epirubicin | 150.0076 |
| Daunorubicin | Sunitinib | 150.0026 |
| Paclitaxel | Afatinib | 149.6683 |
| Vandetanib | Ciclosporin | 149.6566 |
| Erlotinib | Pentamidine | 149.4911 |
| Docetaxel | Vandetanib | 149.3667 |
| Lapatinib | Tirofiban | 149.3435 |
| Sorafenib | Docetaxel | 149.1686 |
| Vemurafenib | Topotecan | 149.114 |
| Pazopanib | Vinblastine | 149.0504 |
| Vorinostat | Lenalidomide | 148.9689 |
| Sorafenib | Hydrocortisone | 148.8624 |
| Hydrocortisone | Gefitinib | 148.7325 |
| Topotecan | Imatinib | 148.6783 |
| Rosuvastatin | Pentamidine | 148.5066 |
| Bosutinib | Epirubicin | 148.4238 |
| Topotecan | Pazopanib | 148.3029 |
| Simvastatin | Imatinib | 148.1823 |
| Paclitaxel | Vandetanib | 148.0012 |
| Paclitaxel | Sunitinib | 147.9866 |
| Daunorubicin | Dasatinib | 147.9826 |
| Lapatinib | Lenalidomide | 147.9195 |
| Daunorubicin | Afatinib | 147.8914 |
| Paclitaxel | Fludarabine | 147.7396 |
| Epirubicin | Dasatinib | 147.6393 |
| Vorinostat | Docetaxel | 147.5857 |
| Vorinostat | Ezetimibe | 147.466 |
| Ceftriaxone | Pentamidine | 147.4081 |
| Doxorubicin | Docetaxel | 147.0707 |
| Lapatinib | Daunorubicin | 147.0521 |
| Vorinostat | Rosuvastatin | 147.043 |
| Pazopanib | Docetaxel | 147.0221 |
| Lapatinib | Montelukast | 146.8397 |
| Nilotinib | Doxorubicin | 146.7087 |
| Simvastatin | Gefitinib | 146.5636 |
| Simvastatin | Pentamidine | 146.4769 |
| Simvastatin | Erlotinib | 146.4032 |
| Vemurafenib | Irinotecan | 146.3449 |
| Pentamidine | Diltiazem | 146.011 |
| Vorinostat | Etoposide | 145.9996 |
| Bortezomib | Pentamidine | 145.9326 |
| Paclitaxel | Erlotinib | 145.9032 |
| Bosutinib | Doxorubicin | 145.8796 |
| Bortezomib | Erlotinib | 145.7559 |
| Bosutinib | Pentamidine | 145.5648 |
| Hydrocortisone | Vandetanib | 145.2811 |
| Vorinostat | Dasatinib | 145.2458 |
| Sunitinib | Tirofiban | 145.1726 |
| Ceftriaxone | Daunorubicin | 145.1301 |
| Topotecan | Daunorubicin | 145.0689 |
| Doxorubicin | Dasatinib | 144.7879 |
| Erlotinib | Etoposide | 144.5884 |
| Bortezomib | Dasatinib | 144.5413 |
| Lapatinib | Rosuvastatin | 144.5013 |
| Simvastatin | Fludarabine | 144.4978 |
| Docetaxel | Pentamidine | 144.3835 |
| Lapatinib | Fludarabine | 144.1955 |
| Vorinostat | Hydrocortisone | 144.1207 |
| Vorinostat | Ciprofloxacin | 143.7832 |
| Docetaxel | Gefitinib | 143.7772 |
| Prednisolone | Dasatinib | 143.7727 |
| Simvastatin | Afatinib | 143.7111 |
| Topotecan | Cyanocobalamin | 143.7028 |
| Bortezomib | Sunitinib | 143.4868 |
| Sorafenib | Etoposide | 143.3978 |
| Ceftriaxone | Afatinib | 143.3813 |
| Vemurafenib | Imatinib | 143.3036 |
| Vorinostat | Cyanocobalamin | 143.1987 |
| Doxorubicin | Rosuvastatin | 143.0575 |
| Topotecan | Gefitinib | 143.037 |
| Bosutinib | Fludarabine | 142.9348 |
| Vorinostat | Epirubicin | 142.9121 |
| Paclitaxel | Pazopanib | 142.8564 |
| Lapatinib | Diltiazem | 142.8375 |
| Etoposide | Imatinib | 142.6565 |
| Warfarin | Gefitinib | 142.6485 |
| Nilotinib | Hydrocortisone | 142.4773 |
| Sorafenib | Daunorubicin | 142.3549 |
| Vorinostat | Gefitinib | 142.2896 |
| Atorvastatin | Pentamidine | 142.1123 |
| Vorinostat | Paclitaxel | 141.9947 |
| Pentamidine | Levofloxacin | 141.8642 |
| Bortezomib | Warfarin | 141.6903 |
| Gemcitabine | Ciclosporin | 141.6069 |
| Doxorubicin | Ceftriaxone | 141.5534 |
| Bortezomib | Atorvastatin | 141.4732 |
| Dasatinib | Pentamidine | 141.3099 |
| Doxorubicin | Sunitinib | 141.2927 |
| Erlotinib | Hydrocortisone | 141.2497 |
| Sunitinib | Warfarin | 141.1208 |
| Lapatinib | Afatinib | 140.8527 |
| Paclitaxel | Daunorubicin | 140.837 |
| Sirolimus | Docetaxel | 140.799 |
| Topotecan | Hydrocortisone | 140.7921 |
| Bortezomib | Levofloxacin | 140.7595 |
| Rosuvastatin | Afatinib | 140.7539 |
| Sunitinib | Vinblastine | 140.6195 |
| Fludarabine | Hydrocortisone | 140.5713 |
| Topotecan | Docetaxel | 140.2396 |
| Erlotinib | Tirofiban | 140.1524 |
| Erlotinib | Docetaxel | 140.1433 |
| Vorinostat | Atorvastatin | 139.9105 |
| Vorinostat | Ceftriaxone | 139.8785 |
| Pazopanib | Diltiazem | 139.7108 |
| Bortezomib | Montelukast | 139.6807 |
| Celecoxib | Topotecan | 139.6651 |
| Bortezomib | Rosuvastatin | 139.6574 |
| Bortezomib | Fludarabine | 139.6442 |
| Topotecan | Dasatinib | 139.6273 |
| Epirubicin | Gefitinib | 139.5292 |
| Doxorubicin | Azelastine | 139.51 |
| Vemurafenib | Pentamidine | 139.4751 |
| Bosutinib | Cyanocobalamin | 139.4282 |
| Nilotinib | Topotecan | 139.4261 |
| Ceftriaxone | Gefitinib | 139.1942 |
| Atorvastatin | Pazopanib | 139.1114 |
| Sorafenib | Tirofiban | 138.9965 |
| Vandetanib | Etoposide | 138.9691 |
| Gemcitabine | Epirubicin | 138.7943 |
| Lapatinib | Bosutinib | 138.5839 |
| Lapatinib | Ceftriaxone | 138.4392 |
| Vemurafenib | Gefitinib | 138.3663 |
| Lapatinib | Vemurafenib | 138.3092 |
| Lapatinib | Atorvastatin | 138.2125 |
| Topotecan | Fludarabine | 138.1732 |
| Nilotinib | Daunorubicin | 138.1269 |
| Daunorubicin | Vandetanib | 137.8452 |
| Vemurafenib | Ceftriaxone | 137.8109 |
| Sorafenib | Paclitaxel | 137.7453 |
| Epirubicin | Ciclosporin | 137.728 |
| Fludarabine | Erlotinib | 137.6136 |
| Sorafenib | Pentamidine | 137.4149 |
| Paclitaxel | Imatinib | 137.2642 |
| Vemurafenib | Fludarabine | 137.2471 |
| Vorinostat | Sorafenib | 137.1436 |
| Topotecan | Ciclosporin | 136.949 |
| Rosuvastatin | Dasatinib | 136.9014 |
| Pazopanib | Etoposide | 136.858 |
| Vorinostat | Daunorubicin | 136.7054 |
| Epirubicin | Pentamidine | 136.58 |
| Docetaxel | Raloxifene | 136.5508 |
| Fludarabine | Etoposide | 136.4807 |
| Vorinostat | Montelukast | 136.4194 |
| Bosutinib | Montelukast | 136.3974 |
| Doxorubicin | Gefitinib | 136.339 |
| Rosuvastatin | Gefitinib | 136.3143 |
| Nilotinib | Docetaxel | 136.1599 |
| Docetaxel | Cyanocobalamin | 135.921 |
| Simvastatin | Vandetanib | 135.9056 |
| Bosutinib | Warfarin | 135.9025 |
| Bortezomib | Ciprofloxacin | 135.8449 |
| Doxorubicin | Lenalidomide | 135.7788 |
| Pentamidine | Gefitinib | 135.7551 |
| Epirubicin | Afatinib | 135.7216 |
| Sunitinib | Rosuvastatin | 135.7117 |
| Gemcitabine | Etoposide | 135.6148 |
| Tirofiban | Imatinib | 135.4749 |
| Simvastatin | Nilotinib | 135.4685 |
| Bortezomib | Irinotecan | 135.4604 |
| Vorinostat | Warfarin | 135.4024 |
| Bortezomib | Bupivacaine | 135.3545 |
| Sirolimus | Etoposide | 135.2765 |
| Etoposide | Ciclosporin | 135.0863 |
| Doxorubicin | Afatinib | 135.016 |
| Afatinib | Lovastatin | 134.9922 |
| Rosuvastatin | Vinblastine | 134.6537 |
| Vemurafenib | Lenalidomide | 134.6315 |
| Vemurafenib | Afatinib | 134.5578 |
| Erlotinib | Epirubicin | 134.4647 |
| Erlotinib | Rosuvastatin | 134.3574 |
| Ceftriaxone | Vandetanib | 134.3439 |
| Docetaxel | Ciclosporin | 134.3324 |
| Bortezomib | Pantoprazole | 134.3267 |
| Daunorubicin | Pantoprazole | 134.1523 |
| Lenalidomide | Pazopanib | 134.0656 |
| Bortezomib | Cyanocobalamin | 134.0412 |
| Rosuvastatin | Vandetanib | 133.9385 |
| Daunorubicin | Imatinib | 133.7936 |
| Bosutinib | Gemcitabine | 133.7817 |
| Ceftriaxone | Erlotinib | 133.6802 |
| Montelukast | Afatinib | 133.6184 |
| Etoposide | Cyanocobalamin | 133.6101 |
| Topotecan | Rosuvastatin | 133.5348 |
| Daunorubicin | Gefitinib | 133.5206 |
| Bosutinib | Irinotecan | 133.4916 |
| Vorinostat | Raloxifene | 133.3764 |
| Fludarabine | Sunitinib | 133.3496 |
| Fludarabine | Daunorubicin | 133.2192 |
| Pazopanib | Irinotecan | 133.1012 |
| Prednisolone | Gefitinib | 132.9101 |
| Daunorubicin | Erlotinib | 132.846 |
| Docetaxel | Tirofiban | 132.811 |
| Montelukast | Gefitinib | 132.7714 |
| Afatinib | Pentamidine | 132.6858 |
| Bosutinib | Diltiazem | 132.4403 |
| Ciprofloxacin | Pentamidine | 132.4316 |
| Lapatinib | Irinotecan | 132.257 |
| Paclitaxel | Tirofiban | 132.2156 |
| Sunitinib | Quinidine | 132.151 |
| Afatinib | Warfarin | 131.8604 |
| Daunorubicin | Hydrocortisone | 131.7946 |
| Dasatinib | Vinblastine | 131.7034 |
| Lenalidomide | Erlotinib | 131.6812 |
| Fludarabine | Montelukast | 131.6489 |
| Nilotinib | Tirofiban | 131.5838 |
| Doxorubicin | Pantoprazole | 131.5837 |
| Bortezomib | Ceftriaxone | 131.4139 |
| Nilotinib | Prednisolone | 131.3036 |
| Vorinostat | Topotecan | 131.2603 |
| Erlotinib | Vinblastine | 131.2266 |
| Nilotinib | Paclitaxel | 131.2209 |
| Nilotinib | Etoposide | 131.1251 |
| Bortezomib | Sorafenib | 131.0977 |
| Vorinostat | Irinotecan | 131.0388 |
| Bortezomib | Raloxifene | 130.8872 |
| Sorafenib | Lenalidomide | 130.8145 |
| Vemurafenib | Warfarin | 130.7686 |
| Bortezomib | Primaquine | 130.6509 |
| Vemurafenib | Vinblastine | 130.5066 |
| Doxorubicin | Erlotinib | 130.1857 |
| Doxorubicin | Vandetanib | 130.0596 |
| Afatinib | Cyanocobalamin | 129.9745 |
| Vorinostat | Fludarabine | 129.9402 |
| Tirofiban | Dasatinib | 129.9132 |
| Doxorubicin | Pentamidine | 129.7741 |
| Simvastatin | Vorinostat | 129.716 |
| Rosuvastatin | Epirubicin | 129.7134 |
| Sorafenib | Irinotecan | 129.5361 |
| Topotecan | Gemcitabine | 129.503 |
| Lapatinib | Primaquine | 129.3266 |
| Lenalidomide | Docetaxel | 129.292 |
| Prednisolone | Pazopanib | 129.2872 |
| Bosutinib | Lenalidomide | 129.1921 |
| Sorafenib | Quinidine | 129.1697 |
| Pentamidine | Imatinib | 129.0675 |
| Vemurafenib | Prednisolone | 128.9228 |
| Prednisolone | Sunitinib | 128.9005 |
| Simvastatin | Bosutinib | 128.8923 |
| Paclitaxel | Pentamidine | 128.8406 |
| Vorinostat | Afatinib | 128.7853 |
| Montelukast | Dasatinib | 128.7614 |
| Daunorubicin | Pentamidine | 128.5603 |
| Montelukast | Pentamidine | 128.491 |
| Vorinostat | Pantoprazole | 128.4677 |
| Fludarabine | Epirubicin | 128.3455 |
| Vemurafenib | Dasatinib | 128.3124 |
| Lapatinib | Pantoprazole | 128.2695 |
| Erlotinib | Irinotecan | 128.197 |
| Pazopanib | Warfarin | 128.1895 |
| Vemurafenib | Raloxifene | 128.1762 |
| Sorafenib | Cocaine | 127.8227 |
| Epirubicin | Vandetanib | 127.8013 |
| Ceftriaxone | Tirofiban | 127.2239 |
| Lapatinib | Furosemide | 127.0856 |
| Sunitinib | Montelukast | 127.0397 |
| Ceftriaxone | Dasatinib | 126.9957 |
| Daunorubicin | Warfarin | 126.9807 |
| Daunorubicin | Atorvastatin | 126.9469 |
| Raloxifene | Ciclosporin | 126.938 |
| Bortezomib | Repaglinide | 126.9155 |
| Ciclosporin | Imatinib | 126.8915 |
| Afatinib | Diltiazem | 126.8501 |
| Vorinostat | Sildenafil | 126.8339 |
| Sorafenib | Fludarabine | 126.785 |
| Probenecid | Sunitinib | 126.7098 |
| Vorinostat | Ofloxacin | 126.5472 |
| Pazopanib | Montelukast | 126.4727 |
| Fludarabine | Warfarin | 126.4312 |
| Epirubicin | Cyanocobalamin | 126.4165 |
| Bosutinib | Pemetrexed | 126.3885 |
| Bortezomib | Chloroquine | 126.3275 |
| Lapatinib | Imatinib | 126.1469 |
| Hydrocortisone | Pentamidine | 125.9467 |
| Vemurafenib | Atorvastatin | 125.8953 |
| Sorafenib | Prednisolone | 125.8435 |
| Primaquine | Sunitinib | 125.816 |
| Lapatinib | Vinblastine | 125.6366 |
| Furosemide | Daunorubicin | 125.6362 |
| Prednisolone | Imatinib | 125.582 |
| Ceftriaxone | Vinblastine | 125.561 |
| Vorinostat | Gemcitabine | 125.5172 |
| Lenalidomide | Etoposide | 125.4848 |
| Ceftriaxone | Sirolimus | 125.4787 |
| Hydrocortisone | Gemcitabine | 125.4246 |
| Furosemide | Gefitinib | 125.407 |
| Warfarin | Imatinib | 125.374 |
| Etoposide | Sildenafil | 125.1431 |
| Daunorubicin | Irinotecan | 125.0956 |
| Tadalafil | Etoposide | 125.0638 |
| Bortezomib | Lenalidomide | 125.0572 |
| Doxorubicin | Imatinib | 124.9581 |
| Vemurafenib | Gemcitabine | 124.9019 |
| Daunorubicin | Montelukast | 124.8094 |
| Tirofiban | Etoposide | 124.6769 |
| Bortezomib | Indometacin | 124.6166 |
| Lapatinib | Gemcitabine | 124.6134 |
| Sorafenib | Montelukast | 124.5747 |
| Bosutinib | Valdecoxib | 124.5435 |
| Fludarabine | Gefitinib | 124.4458 |
| Paclitaxel | Sirolimus | 124.3824 |
| Vemurafenib | Pantoprazole | 124.363 |
| Bortezomib | Vildagliptin | 124.3427 |
| Doxorubicin | Fludarabine | 124.3323 |
| Ceftriaxone | Sunitinib | 124.3221 |
| Lenalidomide | Gefitinib | 124.2743 |
| Fludarabine | Dasatinib | 124.236 |
| Sunitinib | Irinotecan | 124.1734 |
| Doxorubicin | Sorafenib | 124.0798 |
| Bosutinib | Vemurafenib | 124.0178 |
| Daunorubicin | Tirofiban | 123.9685 |
| Pentamidine | Warfarin | 123.8269 |
| Doxorubicin | Gemcitabine | 123.7234 |
| Lapatinib | Pemetrexed | 123.63 |
| Doxorubicin | Hydrocortisone | 123.6205 |
| Simvastatin | Doxorubicin | 123.395 |
| Nilotinib | Vinblastine | 123.2983 |
| Pazopanib | Levofloxacin | 123.2862 |
| Bosutinib | Raloxifene | 123.234 |
| Simvastatin | Topotecan | 123.2272 |
| Lenalidomide | Afatinib | 123.2159 |
| Fludarabine | Afatinib | 123.2155 |
| Irinotecan | Afatinib | 123.0901 |
| Fludarabine | Irinotecan | 123.0768 |
| Bortezomib | Probenecid | 122.9121 |
| Vorinostat | Prednisolone | 122.8484 |
| Pioglitazone | Etoposide | 122.7827 |
| Sorafenib | Cyanocobalamin | 122.7684 |
| Topotecan | Furosemide | 122.7642 |
| Sirolimus | Raloxifene | 122.4805 |
| Simvastatin | Gemcitabine | 122.384 |
| Pemetrexed | Ciclosporin | 122.3801 |
| Topotecan | Etoposide | 122.3059 |
| Topotecan | Warfarin | 122.2506 |
| Paclitaxel | Topotecan | 122.1733 |
| Daunorubicin | Etoposide | 122.1316 |
| Lapatinib | Vandetanib | 122.0669 |
| Vorinostat | Vinblastine | 122.0087 |
| Fludarabine | Pazopanib | 121.9864 |
| Sorafenib | Bupivacaine | 121.9564 |
| Lapatinib | Cyanocobalamin | 121.9083 |
| Docetaxel | Gemcitabine | 121.7638 |
| Sorafenib | Atorvastatin | 121.6967 |
| Bortezomib | Imatinib | 121.6638 |
| Sorafenib | Rosuvastatin | 121.514 |
| Doxorubicin | Saxagliptin | 121.4681 |
| Lenalidomide | Fludarabine | 121.4049 |
| Sunitinib | Diltiazem | 121.3391 |
| Vemurafenib | Levofloxacin | 121.3237 |
| Sorafenib | Ceftriaxone | 121.2868 |
| Daunorubicin | Gemcitabine | 121.1911 |
| Pazopanib | Ciprofloxacin | 121.1215 |
| Topotecan | Afatinib | 121.1188 |
| Epirubicin | Sildenafil | 121.1146 |
| Tirofiban | Ciclosporin | 121.0665 |
| Vorinostat | Vemurafenib | 121.0357 |
| Sirolimus | Pantoprazole | 121.0284 |
| Dorzolamide | Ciclosporin | 120.9297 |
| Bortezomib | Vorinostat | 120.8714 |
| Bortezomib | Pemetrexed | 120.8115 |
| Docetaxel | Sildenafil | 120.7991 |
| Lapatinib | Sorafenib | 120.7799 |
| Tirofiban | Gefitinib | 120.673 |
| Fludarabine | Pentamidine | 120.6098 |
| Docetaxel | Pemetrexed | 120.1991 |
| Doxorubicin | Irinotecan | 120.1401 |
| Bosutinib | Prednisolone | 120.1047 |
| Primaquine | Sorafenib | 120.0872 |
| Bortezomib | Saxagliptin | 120.0839 |
| Sorafenib | Ofloxacin | 119.9175 |
| Rosuvastatin | Raloxifene | 119.8916 |
| Dasatinib | Pantoprazole | 119.8603 |
| Bosutinib | Furosemide | 119.8577 |
| Vemurafenib | Lovastatin | 119.8419 |
| Gemcitabine | Warfarin | 119.7562 |
| Furosemide | Sunitinib | 119.716 |
| Sorafenib | Warfarin | 119.6645 |
| Vandetanib | Warfarin | 119.6145 |
| Topotecan | Raloxifene | 119.5737 |
| Vorinostat | Indometacin | 119.5204 |
| Lapatinib | Erlotinib | 119.4168 |
| Fludarabine | Diltiazem | 119.3589 |
| Paclitaxel | Raloxifene | 119.2525 |
| Lapatinib | Bupivacaine | 119.2302 |
| Quinidine | Erlotinib | 119.223 |
| Topotecan | Sildenafil | 119.21 |
| Nilotinib | Diltiazem | 119.2091 |
| Montelukast | Vinblastine | 119.1695 |
| Nilotinib | Pentamidine | 119.1673 |
| Pentamidine | Cyanocobalamin | 119.1267 |
| Pazopanib | Pentamidine | 119.0563 |
| Vinblastine | Ciclosporin | 119.0226 |
| Bortezomib | Pazopanib | 118.8791 |
| Bortezomib | Gemcitabine | 118.8603 |
| Ceftriaxone | Imatinib | 118.8094 |
| Lapatinib | Quinidine | 118.8064 |
| Methotrexate | Docetaxel | 118.7451 |
| Fludarabine | Vandetanib | 118.7376 |
| Pentamidine | Sildenafil | 118.721 |
| Nilotinib | Warfarin | 118.7061 |
| Vandetanib | Pentamidine | 118.6644 |
| Erlotinib | Montelukast | 118.6497 |
| Lapatinib | Ciprofloxacin | 118.6269 |
| Paclitaxel | Ciclosporin | 118.5603 |
| Vandetanib | Cyanocobalamin | 118.4996 |
| Ceftriaxone | Fludarabine | 118.483 |
| Prednisolone | Erlotinib | 118.4614 |
| Gemcitabine | Afatinib | 118.4245 |
| Celecoxib | Daunorubicin | 118.3236 |
| Atorvastatin | Erlotinib | 118.291 |
| Docetaxel | Vinblastine | 118.2417 |
| Etoposide | Glimepiride | 118.2175 |
| Celecoxib | Repaglinide | 118.106 |
| Pazopanib | Cyanocobalamin | 118.0691 |
| Erlotinib | Gemcitabine | 117.9694 |
| Iloprost | Imatinib | 117.9609 |
| Furosemide | Dasatinib | 117.7497 |
| Topotecan | Pentamidine | 117.7053 |
| Raloxifene | Warfarin | 117.6607 |
| Nilotinib | Ezetimibe | 117.6501 |
| Fludarabine | Imatinib | 117.3572 |
| Topotecan | Sirolimus | 117.354 |
| Simvastatin | Irinotecan | 117.3136 |
| Sirolimus | Rosuvastatin | 117.2267 |
| Doxorubicin | Montelukast | 117.2171 |
| Paclitaxel | Lenalidomide | 117.1628 |
| Vemurafenib | Rosuvastatin | 117.0274 |
| Lapatinib | Levofloxacin | 117.0025 |
| Vemurafenib | Repaglinide | 116.9881 |
| Topotecan | Prednisolone | 116.7832 |
| Montelukast | Vandetanib | 116.7715 |
| Doxorubicin | Warfarin | 116.7172 |
| Paclitaxel | Cyanocobalamin | 116.6646 |
| Paclitaxel | Tadalafil | 116.6429 |
| Ofloxacin | Pazopanib | 116.5965 |
| Celecoxib | Vinblastine | 116.5767 |
| Topotecan | Valdecoxib | 116.5421 |
| Montelukast | Imatinib | 116.4853 |
| Nilotinib | Irinotecan | 116.481 |
| Epirubicin | Raloxifene | 116.4318 |
| Bosutinib | Vorinostat | 116.3873 |
| Dipyridamole | Docetaxel | 116.3186 |
| Lenalidomide | Vandetanib | 116.2602 |
| Bupivacaine | Erlotinib | 116.2324 |
| Bosutinib | Indometacin | 116.1521 |
| Iloprost | Ciclosporin | 116.1096 |
| Prednisolone | Fludarabine | 116.0972 |
| Probenecid | Erlotinib | 115.8853 |
| Furosemide | Erlotinib | 115.8664 |
| Lapatinib | Ofloxacin | 115.7579 |
| Vinblastine | Pantoprazole | 115.7474 |
| Lenalidomide | Epirubicin | 115.6418 |
| Irinotecan | Tirofiban | 115.6167 |
| Bortezomib | Furosemide | 115.604 |
| Saxagliptin | Imatinib | 115.534 |
| Paclitaxel | Gemcitabine | 115.4815 |
| Erlotinib | Cyanocobalamin | 115.447 |
| Topotecan | Trimethoprim | 115.3309 |
| Lenalidomide | Sunitinib | 115.3158 |
| Rosuvastatin | Imatinib | 115.2645 |
| Lapatinib | Gefitinib | 115.1909 |
| Topotecan | Indometacin | 115.1844 |
| Lapatinib | Sunitinib | 115.1662 |
| Nilotinib | Vorinostat | 115.1445 |
| Topotecan | Glimepiride | 115.0856 |
| Simvastatin | Sirolimus | 114.9598 |
| Vemurafenib | Sorafenib | 114.8307 |
| Erlotinib | Vildagliptin | 114.8022 |
| Methotrexate | Etoposide | 114.7763 |
| Etoposide | Pentamidine | 114.7708 |
| Erlotinib | Pantoprazole | 114.7422 |
| Doxorubicin | Atorvastatin | 114.734 |
| Doxorubicin | Etoposide | 114.6633 |
| Sorafenib | Levofloxacin | 114.6282 |
| Saxagliptin | Erlotinib | 114.6044 |
| Tadalafil | Docetaxel | 114.5944 |
| Iloprost | Sunitinib | 114.5286 |
| Pantoprazole | Gefitinib | 114.4831 |
| Lapatinib | Methotrexate | 114.451 |
| Lenalidomide | Dasatinib | 114.4428 |
| Topotecan | Methotrexate | 114.393 |
| Methotrexate | Epirubicin | 114.3731 |
| Bosutinib | Celecoxib | 114.3216 |
| Lovastatin | Gefitinib | 114.2369 |
| Hydrocortisone | Irinotecan | 114.1969 |
| Sirolimus | Epirubicin | 114.102 |
| Hydrocortisone | Tirofiban | 114.0453 |
| Simvastatin | Raloxifene | 114.0218 |
| Vorinostat | Tadalafil | 113.9606 |
| Lapatinib | Raloxifene | 113.957 |
| Atorvastatin | Etoposide | 113.9382 |
| Daunorubicin | Saxagliptin | 113.811 |
| Gemcitabine | Pentamidine | 113.8039 |
| Etoposide | Raloxifene | 113.7664 |
| Paclitaxel | Pemetrexed | 113.6639 |
| Topotecan | Montelukast | 113.6578 |
| Sirolimus | Glimepiride | 113.6488 |
| Bosutinib | Sildenafil | 113.5482 |
| Pemetrexed | Etoposide | 113.5042 |
| Sirolimus | Ciclosporin | 113.4401 |
| Bupivacaine | Daunorubicin | 113.4015 |
| Sunitinib | Saxagliptin | 113.39 |
| Prednisolone | Gemcitabine | 113.3537 |
| Prednisolone | Afatinib | 113.3055 |
| Ofloxacin | Sunitinib | 113.2365 |
| Vemurafenib | Pioglitazone | 113.2055 |
| Doxorubicin | Paclitaxel | 113.1426 |
| Sorafenib | Raloxifene | 113.1415 |
| Topotecan | Pioglitazone | 113.0899 |
| Vemurafenib | Cyanocobalamin | 113.0163 |
| Sirolimus | Valdecoxib | 112.9999 |
| Erlotinib | Warfarin | 112.9773 |
| Pazopanib | Raloxifene | 112.9482 |
| Paclitaxel | Sildenafil | 112.8416 |
| Daunorubicin | Epirubicin | 112.7314 |
| Bosutinib | Pantoprazole | 112.7269 |
| Sirolimus | Montelukast | 112.6813 |
| Fludarabine | Rosuvastatin | 112.6808 |
| Pazopanib | Repaglinide | 112.6795 |
| Afatinib | Raloxifene | 112.6485 |
| Sorafenib | Repaglinide | 112.6126 |
| Celecoxib | Docetaxel | 112.5151 |
| Bortezomib | Etodolac | 112.4998 |
| Vemurafenib | Ciprofloxacin | 112.4111 |
| Nilotinib | Lenalidomide | 112.2375 |
| Pioglitazone | Sirolimus | 112.1718 |
| Atorvastatin | Gefitinib | 112.1456 |
| Bosutinib | Pioglitazone | 112.1163 |
| Lapatinib | Pazopanib | 112.075 |
| Nilotinib | Atorvastatin | 111.9443 |
| Lapatinib | Indometacin | 111.7058 |
| Prednisolone | Daunorubicin | 111.6975 |
| Etodolac | Ciclosporin | 111.6529 |
| Atorvastatin | Dasatinib | 111.6048 |
| Sorafenib | Ciprofloxacin | 111.5625 |
| Rosuvastatin | Etoposide | 111.4927 |
| Lapatinib | Iloprost | 111.4886 |
| Montelukast | Pemetrexed | 111.4632 |
| Doxorubicin | Tirofiban | 111.4527 |
| Bortezomib | Sildenafil | 111.4143 |
| Hydrocortisone | Vinblastine | 111.3643 |
| Nilotinib | Montelukast | 111.3463 |
| Gemcitabine | Vandetanib | 111.2991 |
| Simvastatin | Daunorubicin | 111.2171 |
| Ceftriaxone | Ciclosporin | 111.1686 |
| Bosutinib | Imatinib | 111.1007 |
| Topotecan | Pemetrexed | 111.0782 |
| Simvastatin | Pemetrexed | 111.0638 |
| Lapatinib | Saxagliptin | 111.0388 |
| Bosutinib | Glimepiride | 110.9022 |
| Sunitinib | Atorvastatin | 110.869 |
| Azelastine | Pentamidine | 110.7742 |
| Sunitinib | Pantoprazole | 110.7535 |
| Tirofiban | Diltiazem | 110.7433 |
| Prednisolone | Pentamidine | 110.6482 |
| Simvastatin | Dorzolamide | 110.6333 |
| Sirolimus | Dorzolamide | 110.627 |
| Doxorubicin | Furosemide | 110.5431 |
| Tadalafil | Ciclosporin | 110.4469 |
| Vemurafenib | Tadalafil | 110.3849 |
| Sunitinib | Gemcitabine | 110.3676 |
| Ofloxacin | Pentamidine | 110.2696 |
| Celecoxib | Irinotecan | 110.2511 |
| Quinidine | Imatinib | 110.211 |
| Bortezomib | Tadalafil | 110.2102 |
| Tirofiban | Epirubicin | 110.0356 |
| Afatinib | Valdecocix | 109.9884 |
| Lenalidomide | Vinblastine | 109.8722 |
| Doxorubicin | Bupivacaine | 109.8037 |
| Probenecid | Imatinib | 109.6043 |
| Doxorubicin | Probenecid | 109.4435 |
| Bosutinib | Dipyridamole | 109.4166 |
| Tadalafil | Warfarin | 109.1828 |
| Lenalidomide | Gemcitabine | 108.9966 |
| Vildagliptin | Imatinib | 108.9109 |
| Prednisolone | Vandetanib | 108.9046 |
| Doxorubicin | Repaglinide | 108.8974 |
| Iloprost | Atorvastatin | 108.8756 |
| Nilotinib | Bortezomib | 108.7695 |
| Ofloxacin | Erlotinib | 108.611 |
| Gemcitabine | Gefitinib | 108.5825 |
| Montelukast | Gemcitabine | 108.4474 |
| Bortezomib | Cetorelix | 108.395 |
| Doxorubicin | Celecoxib | 108.3354 |
| Simvastatin | Ceftriaxone | 108.3239 |
| Epirubicin | Pantoprazole | 108.2791 |
| Iloprost | Diltiazem | 108.2217 |
| Sirolimus | Cyanocobalamin | 108.1839 |
| Bupivacaine | Sunitinib | 108.1771 |
| Erlotinib | Repaglinide | 108.0984 |
| Irinotecan | Gemcitabine | 107.9801 |
| Topotecan | Ceftriaxone | 107.9759 |
| Furosemide | Afatinib | 107.7174 |
| Primaquine | Dasatinib | 107.605 |
| Epirubicin | Glimepiride | 107.6003 |
| Lapatinib | Repaglinide | 107.5433 |
| Bosutinib | Ofloxacin | 107.543 |
| Lapatinib | Probenecid | 107.5259 |
| Vemurafenib | Bupivacaine | 107.5032 |
| Pioglitazone | Ciclosporin | 107.4944 |
| Brinzolamide | Ciclosporin | 107.4931 |
| Cyanocobalamin | Gefitinib | 107.438 |
| Doxorubicin | Prednisolone | 107.3794 |
| Vorinostat | Furosemide | 107.3593 |
| Dasatinib | Cyanocobalamin | 107.3324 |
| Bosutinib | Bupivacaine | 107.246 |
| Indometacin | Afatinib | 107.1689 |
| Bosutinib | Etodolac | 107.151 |
| Gemcitabine | Dasatinib | 107.0276 |
| Primaquine | Erlotinib | 107.0172 |
| Rosiglitazone | Etoposide | 106.9985 |
| Lapatinib | Cetorelix | 106.9771 |
| Pioglitazone | Epirubicin | 106.8773 |
| Vemurafenib | Ofloxacin | 106.8406 |
| Doxorubicin | Vildagliptin | 106.7729 |
| Simvastatin | Dipyridamole | 106.7704 |
| Sunitinib | Cyanocobalamin | 106.7276 |
| Ceftriaxone | Etoposide | 106.7243 |
| Pazopanib | Quinidine | 106.7224 |
| Bortezomib | Iloprost | 106.6492 |
| Sorafenib | Chloroquine | 106.5701 |
| Sorafenib | Vildagliptin | 106.5352 |
| Cetorelix | Erlotinib | 106.3875 |
| Repaglinide | Imatinib | 106.2467 |
| Bosutinib | Dorzolamide | 106.1878 |
| Ceftriaxone | Cyanocobalamin | 106.0727 |
| Etodolac | Docetaxel | 106.0647 |
| Celecoxib | Ciclosporin | 106.0263 |
| Furosemide | Vandetanib | 106.0016 |
| Sorafenib | Pantoprazole | 105.9737 |
| Bosutinib | Atorvastatin | 105.9514 |
| Valdecoxib | Ciclosporin | 105.9185 |
| Etoposide | Vinblastine | 105.8607 |
| Azelastine | Fludarabine | 105.8438 |
| Indometacin | Ciclosporin | 105.7954 |
| Ceftriaxone | Iloprost | 105.6551 |
| Sunitinib | Repaglinide | 105.5137 |
| Vorinostat | Repaglinide | 105.4348 |
| Bosutinib | Tadalafil | 105.3139 |
| Sunitinib | Vildagliptin | 105.2802 |
| Bosutinib | Ciprofloxacin | 105.2728 |
| Iloprost | Dasatinib | 105.2694 |
| Irinotecan | Warfarin | 105.24 |
| Azelastine | Ciclosporin | 105.2128 |
| Afatinib | Levofloxacin | 105.2111 |
| Primaquine | Imatinib | 105.1334 |
| Paclitaxel | Dipyridamole | 105.1177 |
| Pazopanib | Rosuvastatin | 105.1121 |
| Dipyridamole | Etoposide | 105.0017 |
| Nilotinib | Primaquine | 104.9952 |
| Doxorubicin | Pemetrexed | 104.9405 |
| Bosutinib | Repaglinide | 104.8515 |
| Pentamidine | Pantoprazole | 104.7288 |
| Bosutinib | Levofloxacin | 104.6401 |
| Furosemide | Pentamidine | 104.5378 |
| Sorafenib | Probenecid | 104.5336 |
| Erlotinib | Raloxifene | 104.4465 |
| Vorinostat | Methotrexate | 104.3016 |
| Sunitinib | Chloroquine | 104.1895 |
| Lapatinib | Chloroquine | 104.1557 |
| Daunorubicin | Repaglinide | 103.9821 |
| Ceftriaxone | Raloxifene | 103.9728 |
| Montelukast | Ciclosporin | 103.8193 |
| Docetaxel | Indometacin | 103.7666 |
| Celecoxib | Afatinib | 103.7621 |
| Celecoxib | Etoposide | 103.7543 |
| Nilotinib | Fludarabine | 103.7305 |
| Nilotinib | Quinidine | 103.6471 |
| Pioglitazone | Docetaxel | 103.642 |
| Bupivacaine | Pazopanib | 103.5861 |
| Cyanocobalamin | Ciclosporin | 103.5756 |
| Repaglinide | Warfarin | 103.5615 |
| Hydrocortisone | Pemetrexed | 103.4778 |
| Sorafenib | Sildenafil | 103.3036 |
| Azelastine | Imatinib | 103.2926 |
| Montelukast | Tirofiban | 103.235 |
| Celecoxib | Quinidine | 103.0787 |
| Glimepiride | Ciclosporin | 102.9617 |
| Celecoxib | Vildagliptin | 102.804 |
| Afatinib | Pantoprazole | 102.7684 |
| Fludarabine | Vinblastine | 102.5501 |
| Tadalafil | Pazopanib | 102.4357 |
| Lapatinib | Vildagliptin | 102.3413 |
| Etodolac | Pantoprazole | 102.3117 |
| Lenalidomide | Iloprost | 102.1982 |
| Nilotinib | Rosuvastatin | 102.1818 |
| Sorafenib | Saxagliptin | 102.0468 |
| Lapatinib | Etodolac | 101.9826 |
| Ciprofloxacin | Afatinib | 101.9553 |
| Pioglitazone | Warfarin | 101.8873 |
| Sunitinib | Raloxifene | 101.8774 |
| Paclitaxel | Rosuvastatin | 101.7957 |
| Pioglitazone | Pentamidine | 101.7362 |
| Erlotinib | Chloroquine | 101.6706 |
| Daunorubicin | Sildenafil | 101.6516 |
| Daunorubicin | Raloxifene | 101.6019 |
| Ceftriaxone | Hydrocortisone | 101.5599 |
| Pentamidine | Lovastatin | 101.4332 |
| Dipyridamole | Ciclosporin | 101.4049 |
| Atorvastatin | Epirubicin | 101.274 |
| Fludarabine | Raloxifene | 101.2557 |
| Lapatinib | Tadalafil | 101.1204 |
| Ceftriaxone | Pazopanib | 101.0657 |
| Dipyridamole | Afatinib | 101.0232 |
| Ofloxacin | Fludarabine | 100.989 |
| Sirolimus | Pemetrexed | 100.8892 |
| Epirubicin | Trimethoprim | 100.8504 |
| Sorafenib | Furosemide | 100.8253 |
| Pazopanib | Indometacin | 100.7777 |
| Daunorubicin | Vildagliptin | 100.7519 |
| Ofloxacin | Imatinib | 100.5585 |
| Sunitinib | Cetorelix | 100.4838 |
| Epirubicin | Valdecocib | 100.4404 |
| Doxorubicin | Dorzolamide | 100.3058 |
| Lapatinib | Pioglitazone | 100.3008 |
| Pentamidine | Repaglinide | 100.2025 |
| Tadalafil | Epirubicin | 100.1508 |
| Bosutinib | Trimethoprim | 100.1291 |
| Etodolac | Rosuvastatin | 100.0802 |
| Daunorubicin | Ciprofloxacin | 99.90386 |
| Nilotinib | Cyanocobalamin | 99.7714 |
| Tadalafil | Pentamidine | 99.53594 |
| Daunorubicin | Chloroquine | 99.44891 |
| Afatinib | Glimepiride | 99.39657 |
| Afatinib | Sildenafil | 99.06484 |
| Fludarabine | Levofloxacin | 98.97963 |
| Primaquine | Daunorubicin | 98.94208 |
| Hydrocortisone | Raloxifene | 98.91298 |
| Ceftriaxone | Etodolac | 98.70375 |
| Tadalafil | Sirolimus | 98.64686 |
| Vandetanib | Valdecocib | 98.63819 |
| Fludarabine | Ciprofloxacin | 98.50272 |
| Pioglitazone | Afatinib | 98.42861 |
| Sorafenib | Cetorelix | 98.35419 |
| Quinidine | Dasatinib | 98.35374 |
| Sirolimus | Sildenafil | 98.24823 |
| Sirolimus | Tirofiban | 98.2138 |
| Methotrexate | Pazopanib | 98.18023 |
| Bortezomib | Dobutamine | 98.08591 |
| Raloxifene | Pentamidine | 98.03313 |
| Pentamidine | Glimepiride | 97.98111 |
| Lapatinib | Trimethoprim | 97.76064 |
| Glimepiride | Warfarin | 97.70893 |
| Simvastatin | Brinzolamide | 97.44275 |
| Pazopanib | Etodolac | 97.36843 |
| Sunitinib | Ciprofloxacin | 97.28712 |
| Etoposide | Dorzolamide | 97.10193 |
| Nilotinib | Levofloxacin | 97.08248 |
| Doxorubicin | Chloroquine | 96.38751 |
| Topotecan | Etodolac | 96.22604 |
| Iloprost | Levofloxacin | 95.79137 |
| Sirolimus | Trimethoprim | 95.75017 |
| Tadalafil | Hydrocortisone | 95.64747 |
| Bortezomib | Lomefloxacin | 95.6038 |
| Celecoxib | Saxagliptin | 95.53344 |
| Sirolimus | Dipyridamole | 95.17211 |
| Topotecan | Glyburide | 94.32485 |
| Atorvastatin | Dorzolamide | 93.72612 |
| Rosiglitazone | Warfarin | 92.98989 |
| Fludarabine | Furosemide | 92.23569 |
| Furosemide | Raloxifene | 90.86216 |

**Supplementary Table 4:** Drug combinations predicted by FSM (hypertension data set)

| Drug A | Drug B | Score |
| --- | --- | --- |
| Hydrochlorothiazide | Pyridoxine | 130.1677 |
| Hydrochlorothiazide | Nebivolol | 123.9608 |
| Indapamide | Metoprolol | 118.7325 |
| Hydrochlorothiazide | Labetalol | 116.6919 |
| Indapamide | Atenolol | 116.254 |
| Hydrochlorothiazide | Nadolol | 115.1128 |
| Hydrochlorothiazide | Captopril | 113.8526 |
| Bendroflumethiazide | Metoprolol | 113.3713 |
| Amlodipine | Indapamide | 109.8225 |
| Hydrochlorothiazide | Bisoprolol | 108.9142 |
| Hydrochlorothiazide | Carvedilol | 107.7682 |
| Hydrochlorothiazide | Atenolol | 107.2169 |
| Bendroflumethiazide | Bisoprolol | 106.4596 |
| Hydrochlorothiazide | Zoledronic | 105.6897 |
| Hydrochlorothiazide | Ezetimibe | 105.6058 |
| Hydrochlorothiazide | Cyanocobalamin | 105.4782 |
| Indapamide | Carvedilol | 105.3952 |
| Indapamide | Bisoprolol | 105.372 |
| Amlodipine | Pyridoxine | 104.4939 |
| Bumetanide | Zoledronic | 102.3331 |
| Indapamide | Salbutamol | 102.1586 |
| Indapamide | Zoledronic | 101.7702 |
| Bendroflumethiazide | Zoledronic | 101.0606 |
| Hydrochlorothiazide | Verapamil | 100.9216 |
| Bendroflumethiazide | Carvedilol | 100.1762 |
| Bendroflumethiazide | Atenolol | 99.97079 |
| Amlodipine | Zoledronic | 99.86007 |
| Hydrochlorothiazide | Timolol | 99.76195 |
| Hydrochlorothiazide | Felodipine | 97.76533 |
| Hydrochlorothiazide | Niacin | 97.64281 |
| Furosemide | Pyridoxine | 97.19322 |
| Carvedilol | Zoledronic | 97.12854 |
| Amlodipine | Bendroflumethiazide | 97.02981 |
| Amlodipine | Captopril | 96.68922 |
| Nebivolol | Zoledronic | 96.32316 |
| Hydrochlorothiazide | Metoprolol | 96.02976 |
| Niacin | Carvedilol | 95.27269 |
| Amlodipine | Nebivolol | 95.07134 |
| Amlodipine | Ezetimibe | 95.02103 |
| Amlodipine | Cyanocobalamin | 94.36521 |
| Hydrochlorothiazide | Diltiazem | 93.78482 |
| Furosemide | Nebivolol | 93.64376 |
| Hydrochlorothiazide | Telmisartan | 93.47731 |
| Hydrochlorothiazide | Propranolol | 93.11043 |
| Hydrochlorothiazide | Irbesartan | 93.05561 |
| Bendroflumethiazide | Nebivolol | 92.92365 |
| Labetalol | Zoledronic | 92.75394 |
| Hydrochlorothiazide | Formoterol | 91.90957 |
| Hydrochlorothiazide | Chlorhexidine | 91.7907 |
| Hydrochlorothiazide | Metoclopramide | 91.66165 |
| Amlodipine | Niacin | 91.63958 |
| Amlodipine | Valdecoxib | 91.50321 |
| Captopril | Zoledronic | 91.30375 |
| Nifedipine | Metoprolol | 90.82458 |
| Bisoprolol | Zoledronic | 90.7224 |
| Hydrochlorothiazide | Hydrocortisone | 90.60838 |
| Hydrochlorothiazide | Salmeterol | 90.4122 |
| Hydrochlorothiazide | Indometacin | 90.2518 |
| Metoprolol | Zoledronic | 90.12324 |
| Bendroflumethiazide | Lisinopril | 89.94149 |
| Amlodipine | Timolol | 89.92112 |
| Atenolol | Zoledronic | 89.6406 |
| Atenolol | Pyridoxine | 89.63203 |
| Captopril | Atenolol | 89.53558 |
| Zoledronic | Cyanocobalamin | 89.50895 |
| Felodipine | Atenolol | 89.43009 |
| Captopril | Metoprolol | 89.23359 |
| Hydrochlorothiazide | Estradiol | 88.96108 |
| Hydrochlorothiazide | Lidocaine | 88.80178 |
| Ezetimibe | Zoledronic | 88.3026 |
| Ezetimibe | Bendroflumethiazide | 88.19027 |
| Nifedipine | Carvedilol | 88.13206 |
| Valdecoxib | Metoprolol | 87.99091 |
| Timolol | Zoledronic | 87.55443 |
| Hydrochlorothiazide | Daunorubicin | 87.47179 |
| Hydrochlorothiazide | Salbutamol | 87.21456 |
| Niacin | Diltiazem | 87.04362 |
| Hydrochlorothiazide | Ribavirin | 86.76716 |
| Niacin | Atenolol | 86.70207 |
| Indapamide | Pyridoxine | 86.49587 |
| Bendroflumethiazide | Propranolol | 86.28032 |
| Indapamide | Irbesartan | 86.11065 |
| Indapamide | Propranolol | 86.11064 |
| Indometacin | Zoledronic | 85.83342 |
| Indapamide | Salmeterol | 85.80604 |
| Acetaminophen | Timolol | 85.64782 |
| Pyridoxine | Metoprolol | 85.58821 |
| Bendroflumethiazide | Salbutamol | 85.46384 |
| Indapamide | Prednisolone | 85.31642 |
| Furosemide | Labetalol | 85.30912 |
| Hydrochlorothiazide | Doxorubicin | 85.19369 |
| Bendroflumethiazide | Hydrocortisone | 85.06523 |
| Atenolol | Bumetanide | 85.00232 |
| Hydrochlorothiazide | Nifedipine | 84.72426 |
| Hydrochlorothiazide | Amlodipine | 84.70875 |
| Pyridoxine | Zoledronic | 84.56384 |
| Hydrochlorothiazide | Montelukast | 84.38509 |
| Indapamide | Lisinopril | 84.25243 |
| Hydrochlorothiazide | Pioglitazone | 84.17904 |
| Amlodipine | Bumetanide | 84.14537 |
| Indapamide | Diltiazem | 84.07232 |
| Captopril | Salbutamol | 84.04992 |
| Verapamil | Zoledronic | 84.04574 |
| Propranolol | Zoledronic | 84.01698 |
| Amlodipine | Carvedilol | 83.96182 |
| Indapamide | Verapamil | 83.7561 |
| Bumetanide | Metoprolol | 83.66091 |
| Topiramate | Timolol | 83.39821 |
| Nadolol | Zoledronic | 83.02163 |
| Niacin | Metoprolol | 83.00261 |
| Furosemide | Carvedilol | 82.93505 |
| Hydrochlorothiazide | Tadalafil | 82.8299 |
| Dexamethasone | Indapamide | 82.76535 |
| Hydrochlorothiazide | Dexamethasone | 82.57526 |
| Pyridoxine | Lisinopril | 82.51367 |
| Dexamethasone | Bendroflumethiazide | 82.42834 |
| Indapamide | Hydrocortisone | 82.40452 |
| Ezetimibe | Atenolol | 82.11447 |
| Diclofenac | Captopril | 81.6681 |
| Amlodipine | Formoterol | 81.6595 |
| Indapamide | Labetalol | 81.64351 |
| Captopril | Rosuvastatin | 81.48271 |
| Hydrochlorothiazide | Diclofenac | 81.41568 |
| Amlodipine | Labetalol | 81.34022 |
| Epinephrine | Zoledronic | 81.27262 |
| Metoprolol | Cyanocobalamin | 81.04147 |
| Formoterol | Zoledronic | 80.89438 |
| Atenolol | Cyanocobalamin | 80.87422 |
| Indapamide | Formoterol | 80.83132 |
| Rosuvastatin | Nebivolol | 80.73266 |
| Loperamide | Indapamide | 80.63299 |
| Methotrexate | Pyridoxine | 80.58114 |
| Captopril | Bisoprolol | 80.48102 |
| Niacin | Cyanocobalamin | 80.33179 |
| Amlodipine | Propranolol | 80.2062 |
| Rosuvastatin | Atenolol | 79.82064 |
| Bendroflumethiazide | Atorvastatin | 79.8133 |
| Acetaminophen | Nebivolol | 79.81007 |
| Estradiol | Indapamide | 79.7058 |
| Hydrochlorothiazide | Lovastatin | 79.67926 |
| Furosemide | Verapamil | 79.6736 |
| Amlodipine | Topiramate | 79.63131 |
| Telmisartan | Furosemide | 79.60345 |
| Nebivolol | Cyanocobalamin | 79.59691 |
| Loperamide | Bendroflumethiazide | 79.54774 |
| Topiramate | Atenolol | 79.3456 |
| Hydrochlorothiazide | Bicalutamide | 79.24348 |
| Bendroflumethiazide | Labetalol | 79.09036 |
| Chlorhexidine | Atenolol | 79.07087 |
| Topiramate | Irbesartan | 78.96642 |
| Bendroflumethiazide | Verapamil | 78.94962 |
| Amlodipine | Chlorhexidine | 78.90852 |
| Diclofenac | Indapamide | 78.88792 |
| Doxorubicin | Indapamide | 78.83532 |
| Irbesartan | Zoledronic | 78.77264 |
| Furosemide | Bisoprolol | 78.73196 |
| Amlodipine | Furosemide | 78.56903 |
| Irbesartan | Pyridoxine | 78.49108 |
| Amlodipine | Ribavirin | 78.26291 |
| Labetalol | Rosuvastatin | 78.24121 |
| Bendroflumethiazide | Pyridoxine | 78.2256 |
| Bendroflumethiazide | Irbesartan | 78.21892 |
| Indapamide | Ezetimibe | 78.09107 |
| Rosuvastatin | Pyridoxine | 78.02238 |
| Niacin | Irbesartan | 77.89969 |
| Hydrochlorothiazide | Glimepiride | 77.8051 |
| Felodipine | Carvedilol | 77.73815 |
| Amlodipine | Nadolol | 77.47172 |
| Diltiazem | Zoledronic | 77.40623 |
| Nifedipine | Nebivolol | 77.34635 |
| Nifedipine | Cyanocobalamin | 77.28955 |
| Amlodipine | Indometacin | 77.07946 |
| Hydrocortisone | Zoledronic | 77.05539 |
| Hydrochlorothiazide | Sildenafil | 77.04853 |
| Acebutolol | Zoledronic | 76.83187 |
| Topiramate | Zoledronic | 76.66577 |
| Amlodipine | Diclofenac | 76.63764 |
| Niacin | Atorvastatin | 76.39969 |
| Furosemide | Captopril | 76.38714 |
| Topiramate | Captopril | 76.37453 |
| Ezetimibe | Metoprolol | 76.35008 |
| Bendroflumethiazide | Diltiazem | 76.34811 |
| Bendroflumethiazide | Captopril | 76.32898 |
| Diclofenac | Carvedilol | 76.31506 |
| Topiramate | Labetalol | 76.28296 |
| Pyridoxine | Bisoprolol | 76.07824 |
| Diclofenac | Cyanocobalamin | 76.03247 |
| Bumetanide | Bisoprolol | 76.01649 |
| Nifedipine | Indapamide | 75.87433 |
| Ezetimibe | Bumetanide | 75.66919 |
| Indapamide | Cyanocobalamin | 75.66562 |
| Ezetimibe | Irbesartan | 75.63361 |
| Topiramate | Nebivolol | 75.63286 |
| Hydrochlorothiazide | Rosuvastatin | 75.56171 |
| Salmeterol | Bendroflumethiazide | 75.54984 |
| Niacin | Bisoprolol | 75.48628 |
| Amlodipine | Sildenafil | 75.46539 |
| Diclofenac | Nebivolol | 75.42935 |
| Chlorhexidine | Metoprolol | 75.30092 |
| Diclofenac | Zoledronic | 75.21373 |
| Hydrochlorothiazide | Atorvastatin | 75.19275 |
| Lisinopril | Zoledronic | 75.17764 |
| Celecoxib | Bisoprolol | 75.15017 |
| Hydrochlorothiazide | Lisinopril | 75.08823 |
| Timolol | Furosemide | 75.0787 |
| Dexamethasone | Captopril | 75.05625 |
| Irbesartan | Furosemide | 75.0166 |
| Indapamide | Telmisartan | 74.7847 |
| Furosemide | Zoledronic | 74.78012 |
| Methotrexate | Felodipine | 74.7396 |
| Indapamide | Nebivolol | 74.64628 |
| Rosuvastatin | Bisoprolol | 74.59824 |
| Indapamide | Timolol | 74.48431 |
| Celecoxib | Atenolol | 74.45714 |
| Bendroflumethiazide | Cyanocobalamin | 74.44852 |
| Telmisartan | Zoledronic | 74.30323 |
| Niacin | Bumetanide | 74.2992 |
| Amlodipine | Celecoxib | 74.28311 |
| Zoledronic | Trimethoprim | 74.27343 |
| Rosuvastatin | Metoprolol | 74.21714 |
| Nifedipine | Timolol | 74.19743 |
| Celecoxib | Pyridoxine | 74.12903 |
| Bumetanide | Carvedilol | 74.11784 |
| Indapamide | Furosemide | 74.07988 |
| Indapamide | Captopril | 74.06173 |
| Valdecoxib | Nebivolol | 74.03482 |
| Diclofenac | Bendroflumethiazide | 74.01455 |
| Captopril | Nebivolol | 74.01126 |
| Bisoprolol | Cyanocobalamin | 74.00814 |
| Hydrochlorothiazide | Trimethoprim | 73.94013 |
| Diclofenac | Irbesartan | 73.92903 |
| Felodipine | Zoledronic | 73.89544 |
| Felodipine | Salbutamol | 73.89216 |
| Bumetanide | Nebivolol | 73.77174 |
| Topiramate | Ezetimibe | 73.62698 |
| Irbesartan | Rosuvastatin | 73.50169 |
| Atorvastatin | Pyridoxine | 73.47551 |
| Captopril | Pyridoxine | 73.37941 |
| Formoterol | Bendroflumethiazide | 73.31783 |
| Topiramate | Metoprolol | 73.25097 |
| Niacin | Furosemide | 73.23984 |
| Diclofenac | Pyridoxine | 73.16749 |
| Niacin | Zoledronic | 73.11682 |
| Furosemide | Felodipine | 73.11359 |
| Bumetanide | Lisinopril | 73.09892 |
| Indapamide | Atorvastatin | 73.06201 |
| Pyridoxine | Diltiazem | 73.00014 |
| Doxorubicin | Niacin | 72.97439 |
| Rosuvastatin | Carvedilol | 72.88987 |
| Felodipine | Lisinopril | 72.82118 |
| Niacin | Timolol | 72.80399 |
| Irbesartan | Bumetanide | 72.73812 |
| Ezetimibe | Salbutamol | 72.71335 |
| Nifedipine | Ezetimibe | 72.7076 |
| Furosemide | Cyanocobalamin | 72.70584 |
| Hydrochlorothiazide | Rosiglitazone | 72.5803 |
| Niacin | Labetalol | 72.53445 |
| Pyridoxine | Carvedilol | 72.40424 |
| Pyridoxine | Cyanocobalamin | 72.38983 |
| Salbutamol | Zoledronic | 72.35052 |
| Captopril | Montelukast | 72.33799 |
| Acetaminophen | Telmisartan | 72.33581 |
| Furosemide | Atenolol | 72.22842 |
| Amlodipine | Bicalutamide | 72.21467 |
| Indapamide | Lidocaine | 72.21063 |
| Diclofenac | Timolol | 72.18656 |
| Rosuvastatin | Verapamil | 71.97196 |
| Nifedipine | Salmeterol | 71.92206 |
| Furosemide | Propranolol | 71.77422 |
| Telmisartan | Niacin | 71.72334 |
| Chlorhexidine | Zoledronic | 71.51032 |
| Estradiol | Bendroflumethiazide | 71.47842 |
| Simvastatin | Bendroflumethiazide | 71.46213 |
| Niacin | Nebivolol | 71.46031 |
| Hydrochlorothiazide | Gemfibrozil | 71.45533 |
| Sildenafil | Zoledronic | 71.42029 |
| Salmeterol | Zoledronic | 71.41565 |
| Ezetimibe | Pyridoxine | 71.36452 |
| Methotrexate | Nebivolol | 71.35074 |
| Lidocaine | Zoledronic | 71.29862 |
| Nifedipine | Atorvastatin | 71.28065 |
| Rosuvastatin | Zoledronic | 71.2245 |
| Ezetimibe | Captopril | 71.1352 |
| Bendroflumethiazide | Lidocaine | 71.10088 |
| Amlodipine | Estradiol | 71.09423 |
| Pioglitazone | Furosemide | 71.00472 |
| Nifedipine | Captopril | 70.92941 |
| Telmisartan | Bendroflumethiazide | 70.86056 |
| Ezetimibe | Timolol | 70.85915 |
| Niacin | Verapamil | 70.82067 |
| Hydrochlorothiazide | Pseudoephedrine | 70.80368 |
| Formoterol | Niacin | 70.71832 |
| Niacin | Sildenafil | 70.69446 |
| Estradiol | Zoledronic | 70.47926 |
| Pseudoephedrine | Diltiazem | 70.47454 |
| Acetaminophen | Felodipine | 70.37265 |
| Furosemide | Metoprolol | 70.33833 |
| Salmeterol | Felodipine | 70.32993 |
| Nebivolol | Lidocaine | 70.29211 |
| Ezetimibe | Cyanocobalamin | 70.2588 |
| Dexamethasone | Irbesartan | 70.14698 |
| Nebivolol | Lisinopril | 70.123 |
| Indapamide | Daunorubicin | 70.07705 |
| Irbesartan | Hydrocortisone | 70.06662 |
| Salmeterol | Captopril | 69.96258 |
| Hydralazine | Metoprolol | 69.88053 |
| Pioglitazone | Bendroflumethiazide | 69.87358 |
| Topiramate | Bisoprolol | 69.72413 |
| Hydralazine | Atenolol | 69.69724 |
| Dexamethasone | Zoledronic | 69.65268 |
| Ezetimibe | Bisoprolol | 69.61412 |
| Celecoxib | Verapamil | 69.60345 |
| Nifedipine | Pyridoxine | 69.48942 |
| Atenolol | Sildenafil | 69.42534 |
| Irbesartan | Cyanocobalamin | 69.40997 |
| Nifedipine | Labetalol | 69.39371 |
| Felodipine | Cyanocobalamin | 69.36894 |
| Ezetimibe | Nebivolol | 69.29992 |
| Captopril | Diltiazem | 69.21998 |
| Ezetimibe | Carvedilol | 69.20322 |
| Topiramate | Telmisartan | 69.1676 |
| Bumetanide | Propranolol | 69.125 |
| Amlodipine | Hydralazine | 68.98057 |
| Niacin | Bendroflumethiazide | 68.96012 |
| Daunorubicin | Zoledronic | 68.84168 |
| Indometacin | Metoprolol | 68.82426 |
| Dexamethasone | Pyridoxine | 68.80582 |
| Chlorhexidine | Bisoprolol | 68.68408 |
| Nifedipine | Bumetanide | 68.61872 |
| Amlodipine | Trimethoprim | 68.54702 |
| Ribavirin | Metoprolol | 68.50477 |
| Diclofenac | Diltiazem | 68.48454 |
| Nifedipine | Bendroflumethiazide | 68.48199 |
| Nifedipine | Zoledronic | 68.41831 |
| Rosuvastatin | Bumetanide | 68.41378 |
| Hydrochlorothiazide | Methotrexate | 68.35325 |
| Acetaminophen | Verapamil | 68.2301 |
| Bendroflumethiazide | Rosuvastatin | 68.16275 |
| Diclofenac | Atenolol | 68.14078 |
| Diclofenac | Ezetimibe | 68.08716 |
| Celecoxib | Ezetimibe | 68.05585 |
| Nebivolol | Diltiazem | 68.03541 |
| Diclofenac | Sildenafil | 68.01061 |
| Doxorubicin | Zoledronic | 67.99741 |
| Hydrochlorothiazide | Tamoxifen | 67.94474 |
| Acetaminophen | Labetalol | 67.87939 |
| Atorvastatin | Zoledronic | 67.75717 |
| Rosuvastatin | Cyanocobalamin | 67.73899 |
| Indapamide | Niacin | 67.71031 |
| Nifedipine | Formoterol | 67.66053 |
| Furosemide | Tadalafil | 67.64606 |
| Topiramate | Verapamil | 67.62644 |
| Simvastatin | Hydrochlorothiazide | 67.61667 |
| Diclofenac | Bumetanide | 67.61226 |
| Atenolol | Lidocaine | 67.60063 |
| Simvastatin | Indapamide | 67.58184 |
| Bumetanide | Pyridoxine | 67.41672 |
| Lidocaine | Metoprolol | 67.33736 |
| Topiramate | Carvedilol | 67.30539 |
| Atorvastatin | Felodipine | 67.30082 |
| Chlorhexidine | Cyanocobalamin | 67.28891 |
| Telmisartan | Rosuvastatin | 67.23733 |
| Atenolol | Diltiazem | 66.95888 |
| Ezetimibe | Verapamil | 66.95193 |
| Diclofenac | Verapamil | 66.92369 |
| Pyridoxine | Nebivolol | 66.89578 |
| Carvedilol | Lidocaine | 66.83964 |
| Loperamide | Rosuvastatin | 66.83394 |
| Topiramate | Felodipine | 66.81885 |
| Atorvastatin | Nebivolol | 66.7393 |
| Formoterol | Captopril | 66.72526 |
| Timolol | Rosuvastatin | 66.71778 |
| Topiramate | Indometacin | 66.657 |
| Nadolol | Rosuvastatin | 66.656 |
| Atorvastatin | Bumetanide | 66.60098 |
| Sildenafil | Metoprolol | 66.5929 |
| Topiramate | Propranolol | 66.50053 |
| Hydrochlorothiazide | Fenofibrate | 66.47354 |
| Methotrexate | Zoledronic | 66.31108 |
| Diclofenac | Labetalol | 66.25015 |
| Captopril | Bumetanide | 66.22974 |
| Celecoxib | Cyanocobalamin | 66.21248 |
| Felodipine | Rosuvastatin | 66.19918 |
| Verapamil | Pyridoxine | 66.17247 |
| Pyridoxine | Sildenafil | 66.17201 |
| Bumetanide | Diltiazem | 65.92781 |
| Diclofenac | Bisoprolol | 65.90868 |
| Nifedipine | Methotrexate | 65.85313 |
| Chlorhexidine | Carvedilol | 65.82843 |
| Celecoxib | Propranolol | 65.79803 |
| Nifedipine | Sildenafil | 65.71489 |
| Indometacin | Bisoprolol | 65.6046 |
| Diclofenac | Telmisartan | 65.59749 |
| Chlorhexidine | Diltiazem | 65.55578 |
| Salbutamol | Valdecoxib | 65.55453 |
| Niacin | Salbutamol | 65.42518 |
| Acetylsalicylic | Timolol | 65.34226 |
| Chlorhexidine | Atorvastatin | 65.26587 |
| Diclofenac | Metoprolol | 65.25006 |
| Topiramate | Lidocaine | 65.17002 |
| Loperamide | Diclofenac | 65.14188 |
| Prednisolone | Irbesartan | 65.13744 |
| Salbutamol | Bumetanide | 65.07375 |
| Hydrochlorothiazide | Theophylline | 65.05921 |
| Carvedilol | Cyanocobalamin | 65.04746 |
| Indapamide | Rosuvastatin | 64.98716 |
| Lidocaine | Bisoprolol | 64.94257 |
| Dexamethasone | Nebivolol | 64.92894 |
| Bendroflumethiazide | Methotrexate | 64.83623 |
| Pyridoxine | Lidocaine | 64.7746 |
| Diltiazem | Bisoprolol | 64.77433 |
| Bicalutamide | Metoprolol | 64.62675 |
| Nebivolol | Sildenafil | 64.61376 |
| Timolol | Pyridoxine | 64.6013 |
| Topiramate | Cyanocobalamin | 64.54876 |
| Ezetimibe | Labetalol | 64.40785 |
| Rosuvastatin | Propranolol | 64.40345 |
| Nifedipine | Furosemide | 64.38796 |
| Acetylsalicylic | Irbesartan | 64.24033 |
| Pyridoxine | Propranolol | 64.19678 |
| Indapamide | Felodipine | 64.189 |
| Prednisolone | Zoledronic | 64.04054 |
| Topiramate | Pyridoxine | 63.8026 |
| Propranolol | Cyanocobalamin | 63.71254 |
| Telmisartan | Ezetimibe | 63.69273 |
| Salmeterol | Niacin | 63.64957 |
| Indapamide | Ribavirin | 63.63544 |
| Nifedipine | Rosuvastatin | 63.58021 |
| Bicalutamide | Carvedilol | 63.50451 |
| Lidocaine | Cyanocobalamin | 63.24429 |
| Lovastatin | Zoledronic | 63.02006 |
| Felodipine | Sildenafil | 63.01841 |
| Estradiol | Irbesartan | 63.00383 |
| Chlorhexidine | Ezetimibe | 62.98775 |
| Montelukast | Zoledronic | 62.90351 |
| Dexamethasone | Ezetimibe | 62.88523 |
| Hydrocortisone | Pyridoxine | 62.86768 |
| Bumetanide | Cyanocobalamin | 62.82404 |
| Doxorubicin | Bendroflumethiazide | 62.75388 |
| Hydralazine | Cyanocobalamin | 62.74141 |
| Hydralazine | Carvedilol | 62.63193 |
| Niacin | Daunorubicin | 62.55272 |
| Loperamide | Bumetanide | 62.4533 |
| Furosemide | Doxazosin | 62.42635 |
| Indapamide | Pioglitazone | 62.26619 |
| Dexamethasone | Timolol | 62.18653 |
| Atorvastatin | Hydralazine | 62.15842 |
| Chlorhexidine | Furosemide | 62.12585 |
| Bumetanide | Lidocaine | 62.0688 |
| Salbutamol | Rosuvastatin | 61.99476 |
| Bicalutamide | Atenolol | 61.92895 |
| Atenolol | Ribavirin | 61.81006 |
| Formoterol | Rosuvastatin | 61.68949 |
| Chlorhexidine | Captopril | 61.6401 |
| Topiramate | Diltiazem | 61.62795 |
| Estradiol | Timolol | 61.57069 |
| Bendroflumethiazide | Glimepiride | 61.53605 |
| Salmeterol | Rosuvastatin | 61.52844 |
| Formoterol | Ezetimibe | 61.47129 |
| Ribavirin | Bisoprolol | 61.2821 |
| Diclofenac | Nadolol | 61.25687 |
| Salbutamol | Pyridoxine | 61.25299 |
| Chlorhexidine | Rosuvastatin | 61.25013 |
| Hydrocortisone | Carvedilol | 61.14661 |
| Furosemide | Ribavirin | 61.06213 |
| Hydrocortisone | Atenolol | 60.97205 |
| Indapamide | Chlorhexidine | 60.80264 |
| Hydralazine | Bisoprolol | 60.79597 |
| Bicalutamide | Zoledronic | 60.77951 |
| Topiramate | Diclofenac | 60.68992 |
| Felodipine | Diltiazem | 60.53226 |
| Formoterol | Furosemide | 60.45959 |
| Chlorhexidine | Verapamil | 60.34291 |
| Sildenafil | Bisoprolol | 60.26162 |
| Topiramate | Atorvastatin | 59.99079 |
| Hydrocortisone | Nebivolol | 59.76353 |
| Salbutamol | Cyanocobalamin | 59.73159 |
| Doxorubicin | Ezetimibe | 59.5038 |
| Captopril | Carvedilol | 59.44989 |
| Telmisartan | Valdecocib | 59.33719 |
| Ezetimibe | Felodipine | 59.3199 |
| Salmeterol | Ezetimibe | 59.16354 |
| Timolol | Hydrocortisone | 59.09772 |
| Topiramate | Rosuvastatin | 58.86543 |
| Salbutamol | Diltiazem | 58.60292 |
| Niacin | Lisinopril | 58.47684 |
| Chlorhexidine | Salbutamol | 58.15871 |
| Salbutamol | Lidocaine | 58.05938 |
| Topiramate | Doxazosin | 57.50324 |
| Ezetimibe | Propranolol | 57.43424 |
| Ribavirin | Bumetanide | 57.22946 |
| Simvastatin | Timolol | 57.10546 |
| Ezetimibe | Daunorubicin | 56.904 |
| Chlorhexidine | Propranolol | 56.85084 |
| Chlorhexidine | Pyridoxine | 56.80238 |
| Diltiazem | Propranolol | 56.55861 |
| Verapamil | Bumetanide | 56.4363 |
| Doxorubicin | Topiramate | 56.38281 |
| Pyridoxine | Pantoprazole | 55.75629 |

